# Global analysis of transcription start sites in the new ovine reference genome (*Oar rambouillet v1.0*)

**DOI:** 10.1101/2020.07.06.189480

**Authors:** Mazdak Salavati, Alex Caulton, Richard Clark, Iveta Gazova, Timothy P. L. Smith, Kim C. Worley, Noelle E. Cockett, Alan L. Archibald, Shannon M. Clarke, Brenda M. Murdoch, Emily L. Clark, on behalf of The Ovine FAANG Project Consortium

## Abstract

The overall aim of the Ovine FAANG project is to provide a comprehensive annotation of the new highly contiguous sheep reference genome sequence (*Oar rambouillet v1.0)*. Mapping of transcription start sites (TSS) is a key first step in understanding transcript regulation and diversity. Using 56 tissue samples collected from the reference ewe Benz2616 we have performed a global analysis of TSS and TSS-Enhancer clusters using Cap Analysis Gene Expression (CAGE) sequencing. CAGE measures RNA expression by 5’ cap-trapping and has been specifically designed to allow the characterization of TSS within promoters to single-nucleotide resolution. We have adapted an analysis pipeline that uses TagDust2 for clean-up and trimming, Bowtie2 for mapping, CAGEfightR for clustering and the Integrative Genomics Viewer (IGV) for visualization. Mapping of CAGE tags indicated that the expression levels of CAGE tag clusters varied across tissues. Expression profiles across tissues were validated using corresponding polyA+ mRNA-Seq data from the same samples. After removal of CAGE tags with < 10 read counts, 39.3% of TSS overlapped with 5’ ends of 31,113 transcripts that had been previously annotated by NCBI (out of a total of 56,308 from the NCBI annotation). For 25,195 of the transcripts, previously annotated by NCBI, no TSS meeting stringent criteria were identified. A further 14.7% of TSS mapped to within 50bp of annotated promoter regions. Intersecting these predicted TSS regions with annotated promoter regions (±50bp) revealed 46% of the predicted TSS were ‘novel’ and previously un-annotated. Using whole genome bisulphite sequencing data from the same tissues we were able to determine that a proportion of these ‘novel’ TSS were hypo-methylated (32.2%) indicating that they are likely to be reproducible rather than ‘noise’. This global analysis of TSS in sheep will significantly enhance the annotation of gene models in the new ovine reference assembly. Our analyses provide one of the highest resolution annotations of transcript regulation and diversity in a livestock species to date.

## Introduction

The Functional Annotation of Animal Genomes (FAANG) consortium is a concerted international effort to use molecular assays, developed during the Human ENCODE project (Birney et al., 2007), to annotate the majority of functional elements in the genomes of domesticated animals (Andersson et al., 2015; Giuffra and Tuggle, 2019). Towards this aim the overarching goal of the Ovine FAANG project (Murdoch, 2019) is to provide a comprehensive annotation of the new highly contiguous reference genome for sheep, *Oar rambouillet v1.0* (https://www.ncbi.nlm.nih.gov/assembly/GCF_002742125.1/). The Ovine FAANG project is developing a deep and robust dataset of expressed elements and regulatory features in the sheep genome as a resource for the livestock genomics community. Here we describe a global analysis of transcription start sites (TSS) using Cap Analysis Gene Expression (CAGE) sequencing.

CAGE measures RNA expression by 5’ cap-trapping to identify the 5’ ends of both polyadenylated and non-polyadenylated RNAs including lncRNAs and miRNAs, and has been specifically designed to allow the characterization of TSS within promoters to single-nucleotide resolution (Takahashi et al., 2012). By using 5’-Cap capture we avoid transcripts that have been 5’ degraded. Conventional RNA-Seq and cDNA datasets can be ‘contaminated’ with such degradation products and data from transcripts where first strand cDNA synthesis was incomplete. These ‘contaminants’ can give rise to erroneous transcript / gene models with false 5’ ends. The level of resolution provided by CAGE allows investigation of the regulatory inputs driving transcript expression, and construction of transcriptional networks to study, for example, the genetic basis for disease susceptibility (Baillie et al., 2017) or for systematic analysis of transcription start sites through development (Lizio et al., 2017). Using CAGE sequencing technology, the FANTOM5 consortium generated a comprehensive annotation of TSS for the human genome, which included the major primary cell and tissue types (Forrest et al., 2014a).

The goal of this study was to annotate TSS and TSS-Enhancer clusters in the ovine genome (*Oar rambouillet v1.0*). Our approach was to perform CAGE analysis on 55 tissues and one type of primary immune cell (alveolar macrophages). Tissues representing all the major organ systems were collected from Benz2616, the Rambouillet ewe used to generate the *Oar rambouillet v1.0* reference assembly. CAGE tags for each tissue sample clustered with a high level of specificity according to their expression profiles as measured by RNA-Seq. Mapping of CAGE tags indicated that a large proportion of detected TSS did not overlap with the current annotated 5’ end of transcripts. The reproducibility of these ‘novel’ TSS was tested using whole genome DNA methylation profiles from a subset of the same tissues.

DNA methylation plays a key role in the regulation of gene expression and the maintenance of genome stability (Ibeagha-Awemu and Zhao, 2015), and is the most highly studied epigenetic mark. In mammalian species, DNA methylation occurs primarily at cytosine-phosphate-guanine dinucleotides (CpG) and to a lesser extent at CHH and CHG sites (where C = Cytosine; H = Adenine, Guanine, or Thymine; and G = Guanine) (An et al., 2018). Generally, DNA methylation in the promoter region of genes represses transcription, inhibiting elongation by transcriptional machinery.

Methylation over TSS blocks transcription initiation; while, conversely, methylation within gene bodies stimulates elongation and influences alternative splicing of transcripts (Jones, 2012; Lev Maor et al., 2015; An et al., 2018). Using DNA methylation profiles, we were able to determine the proportion of ‘novel’ TSS in our dataset that were likely true signals of transcription initiation based on a hypo-methylated state rather than being an artefact of CAGE-sequencing.

We provide the annotation of TSS in the ovine genome as tracks in a genome browser via the Track Hub Registry and visualise these in the R package GViz, ensuring the data is accessible and useable to the livestock genomics community. The global analysis of TSS we present here will significantly enhance the annotation of gene models in the new ovine reference assembly demonstrating the utility of the datasets generated by the Ovine FAANG project and providing a foundation for future work.

## Methods

### Animals

Tissues were collected from an adult female Rambouillet sheep at the Utah Veterinary Diagnostic Laboratory on April 29, 2016. At the time of sample collection Benz2616 was approximately 6 years of age and after a thorough veterinary examination confirmed to be healthy. Benz 2616 was donated to the project by the USDA. Sample collection methods were planned and tested over 15 months in 2015 to 2016, a description of these is available via the FAANG Data Coordination Centre https://data.faang.org/api/fire_api/samples/USU_SOP_Ovine_Benz2616_Tissue_Collection_20160426.pdf .

### Sample collection

Necropsy of Benz2616 was performed by a veterinarian to ensure proper identification of tissues, and a team of scientists on hand provided efficient and rapid transfer of tissue sections to containers which were snap frozen in liquid nitrogen prior to transfer to -80°C for long-term storage. Alveolar macrophages were collected by bronchoalveolar lavage as described in (Cordier et al., 1990). Details of all 100 samples collected from Benz2616 are included in the BioSamples database under submission GSB-7268, group accession number SAMEG329607 (https://www.ebi.ac.uk/biosamples/samples/SAMEG329607) and associated information is recorded according to FAANG metadata specifications (Harrison et al.,

2018). The FAANG assays, as described below, were generated from a subset of tissues for CAGE (56 tissues), polyA+ mRNA-Seq (58 tissues) and WGBS (8 tissues) (Figure 1).

**Figure 1.**
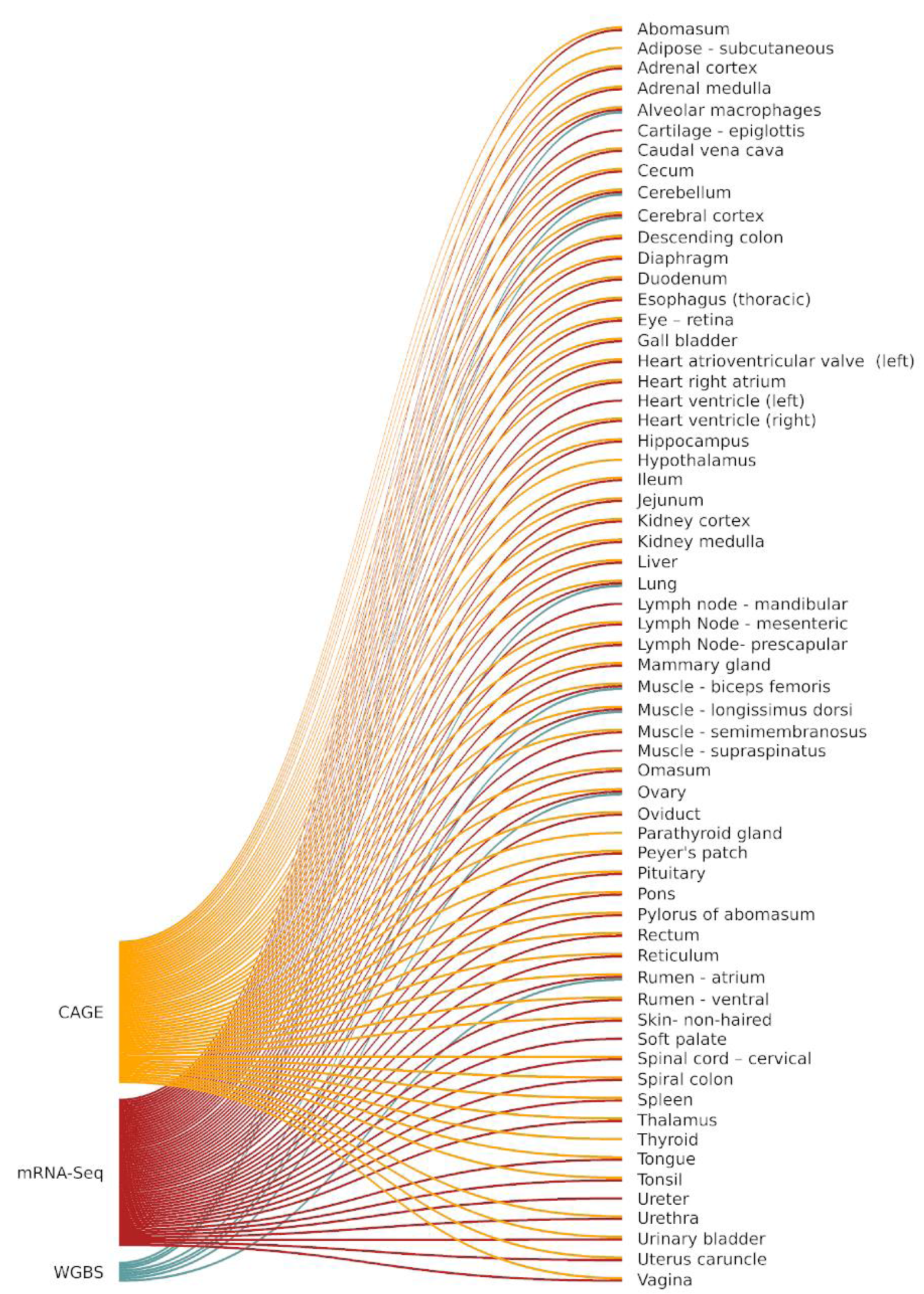
FAANG assays (CAGE, WGBS and mRNA-Seq) performed on each tissue from Benz2616.

### CAGE Library Preparation and Analysis

#### RNA Isolation for CAGE library preparation

Frozen tissues (60-100mg per sample) were homogenised by grinding with a mortar and pestle on dry ice and RNA was isolated using TRIzol Reagent (Invitrogen) according to the manufacturer’s instructions. After RNA isolation 10μg of RNA per sample was treated with DNase I (NEB) then column purified using a RNeasy MinElute kit (Qiagen), according to the manufacturer’s instructions. Full details of the RNA extraction protocol are available via the FAANG Data Coordination Centre https://data.faang.org/api/fire_api/assays/USDA_SOP_RNA_Extraction_Fro <u>m_Tissue_20180626.pdf</u> . Each RNA sample was run on an Agilent BioAnalyzer to ensure RNA integrity was sufficiently high (RIN^e^>6). Details of RNA purity metrics for each sample are included in Supplementary Table 1. RNA samples were then stored at -80°C for downstream analysis.

#### CAGE library preparation and sequencing

CAGE libraries were prepared for each sample as described in (Takahashi et al., 2012) from a starting quantity of 5μg of DNase treated total RNA. Random primers were used to ensure conversion of all 5’ cap-trapping RNAs according to (Takahashi et al., 2012).

The full protocol is available via the FAANG Data Coordination Centre https://data.faang.org/api/fire_api/assays/ROSLIN_SOP_CAGE-library-preparation_20190903.pdf. Libraries were prepared in batches of eight and pooled. Sequencing was performed on the Illumina HiSeq 2500 platform by multiplexing 8 samples on one lane to generate approximately 20 million 50bp single-end reads per sample. Eight of the available fifteen 5’ linker barcodes from (Takahashi et al., 2012) were used for multiplexing: ACG, GAT, CTT, ATG, GTA, GCC, TAG and TGG. In total 8 separate library pools were generated and spread across two HiSeq 2500 flow cells. Details of barcodes assigned to each sample and pool IDs are included in Supplementary Table 1.

#### Processing and mapping of CAGE libraries

All sequence data were processed using in house scripting (bash and R) on the University of Edinburgh high performance computing facility (Edinburgh, 2020). The analysis protocol for CAGE is available via https://data.faang.org/api/fire_api/analysis/ROSLIN_SOP_CAGE_analysis_pipeli <u>ne_20191029.pdf</u> and summarised in Figure 2. To de-multiplex the data we used the FastX toolkit version 0.014 (Hannon Lab, 2017) for short read pre-processing. We then used TagDust2 v.2.33 (Lassmann, 2015) to extract mappable reads from the raw data and for read clean-up to remove the *EcoP1* site and barcode, according to the recommendations of the FANTOM5 consortium e.g. (Bertin et al., 2017). This process resulted in cleaned reads approximately 27nt in length (hereafter referred to as CAGE tags) which were mapped to the Rambouillet Benz2616 genome available from NCBI (*Oar rambouillet v1.0* GCA_002742125.1) using Bowtie2 v.2.3.5.1 in --very-sensitive mode equivalent to options *-D 20 -R 3 -N 0 -L 20 -i S,1,0.50* (Langmead and Salzberg, 2012). Multi-mapped reads were identified using Bowtie2 v.2.3.5.1 in --very-sensitive mode and excluded from the rest of the analysis. The mapped BAM files were then processed for base pair resolution strand specific read counts using bedtools v.2.29.0 (Quinlan and Hall, 2010). Metrics for the attrition of raw reads at each stage of the analysis pipeline are included in Supplementary File 1, Section 1.1. In order for the bedGraph files to be used in the CAGEfightR package they were converted to bigWig format using UCSCs tool BedGraphToBigWig (Kent et al., 2010).

**Figure 2.**
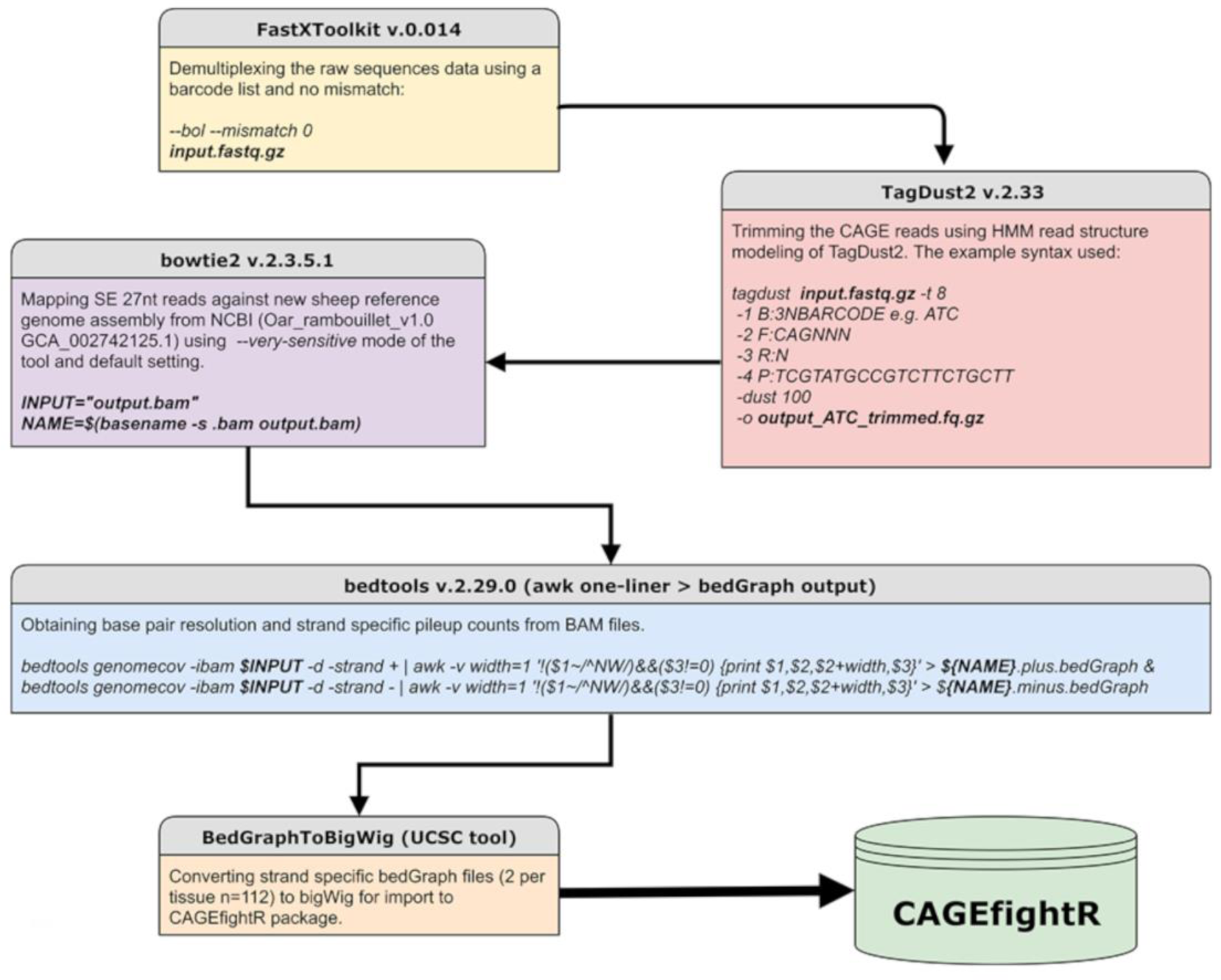
Workflow of the analysis pipeline and respective tools used for CAGE sequence data analysis

#### Normalisation and mapping of CAGE tags

For normalisation and clustering of CAGE tags (as CAGE Tags-Per-Million Mapped: CTPM) we used the software package CAGEfightR v.1.5.1 (Thodberg and Sandelin, 2019). The normalisation was performed by dividing CAGE tag counts in each predicted cluster by the total mapped CAGE tags in the sample, multiplied by 1.0e6. To perform these analyses we created a custom BSgenome object (a container of the genomic sequence) for sheep from *Oar rambouillet v1.0* using the BSgenome Bioconductor package v.1.53.1 (Pages, 2020). Distribution metrics of CAGE tags across the genome were annotated and analysed using the TxDB transcript ID assignment and Genomic Features package v.1.36.4 (Lawrence et al., 2013). The TxDB object was created using the NCBI gff3 gene annotation file from NCBI *Oar rambouillet v1.0* GCA_002742125.1 (*GCF_002742125.1_Oar rambouillet v1.0_genomic.gff release 103*).

#### Clustering of CAGE tags

To annotate TSS in the *Oar rambouillet v1.0* genome assembly we first generated expression read counts for each tag (bp resolution). Tags with < 10 read counts were removed first then any tags that were not present in at least 37/56 tissues (i.e. two thirds of the tissues) were also removed. This conservative representation threshold was introduced to ensure CAGE tags included in downstream analysis were reproducible. In the absence of additional biological replicates, we based this on the assumption that a CAGE tag was more likely to be reproducible if it was shared across multiple tissues. However, it should be noted that this method would reduce sensitivity to putative highly tissue-specific TSS and this is discussed later. Gene annotation from the NCBI’s GTF file was used to validate the coordinates of predicted CAGE clusters (i.e. residing within or outside the promoter of annotated genes). Five thresholds for representation, of CAGE tags (excluding intergenic and intronic tags) across tissues, were compared (1 tissue, 5 tissues, 1/3^rd^ of the tissues, half of the tissues, 2/3^rds^ of the tissues and all of the tissues). The proportion of CAGE tag clusters within (tagged by unique gene IDs) or outside the promoter region (untagged) was used to compare each threshold. Highly stringent filtering (56/56 representation) found CAGE tag clusters associated with 2,949 genes (out of 30,862 genes annotated by NCBI) representing putative TSS for genes expressed in all 56 tissues. A reduction of the threshold to 2/3^rds^ (37/56 tissues) resulted in 13,912 genes (31,113 transcripts) associated with CAGE tag clusters. Reducing the threshold further to 1/3^rd^ of tissues resulted in a high proportion of CAGE tag clusters that were not associated with genes (‘untagged’) (41.6%) and 18,005 associated with genes (39,458 transcripts). According to this criterion we selected the 2/3^rds^ threshold. Although highly stringent this provided only the highest confidence TSS tag clusters, associated with widely expressed genes and widely used promotors, for the analysis of the dataset we present here. Further details of this comparison are included in Supplementary File 1 Section 1.2.

TSS expression profiles (as CTPM) were then regenerated for each tissue using the CAGEfightR v. 1.5.1 quickTSS, quickEnhancers and findLinks functions (Thodberg and Sandelin, 2019). The CAGE tags clustered A) uni-directionally (according to the sense or anti-sense flag of the mapped CAGE tag) into predicted TSS and B) bi-directionally, using the TSS-Enhancer detection algorithm from CAGEfightR (Thodberg and Sandelin, 2019), into correlated TSS and enhancer (TSS-Enhancer) clusters. Bi-directional (TSS-Enhancer) clusters are defined as clusters of CAGE tags that are located on the opposing strand within 400 bp-1000bp proximity of the centre of a promoter (Thodberg and Sandelin, 2019). The bi-directional clusters outside of this range were excluded from this analysis according to the previously described method in (Thodberg et al., 2019). The concept of uni-directional and bi- directional clustering is illustrated in Figure 3.

**Figure 3.**
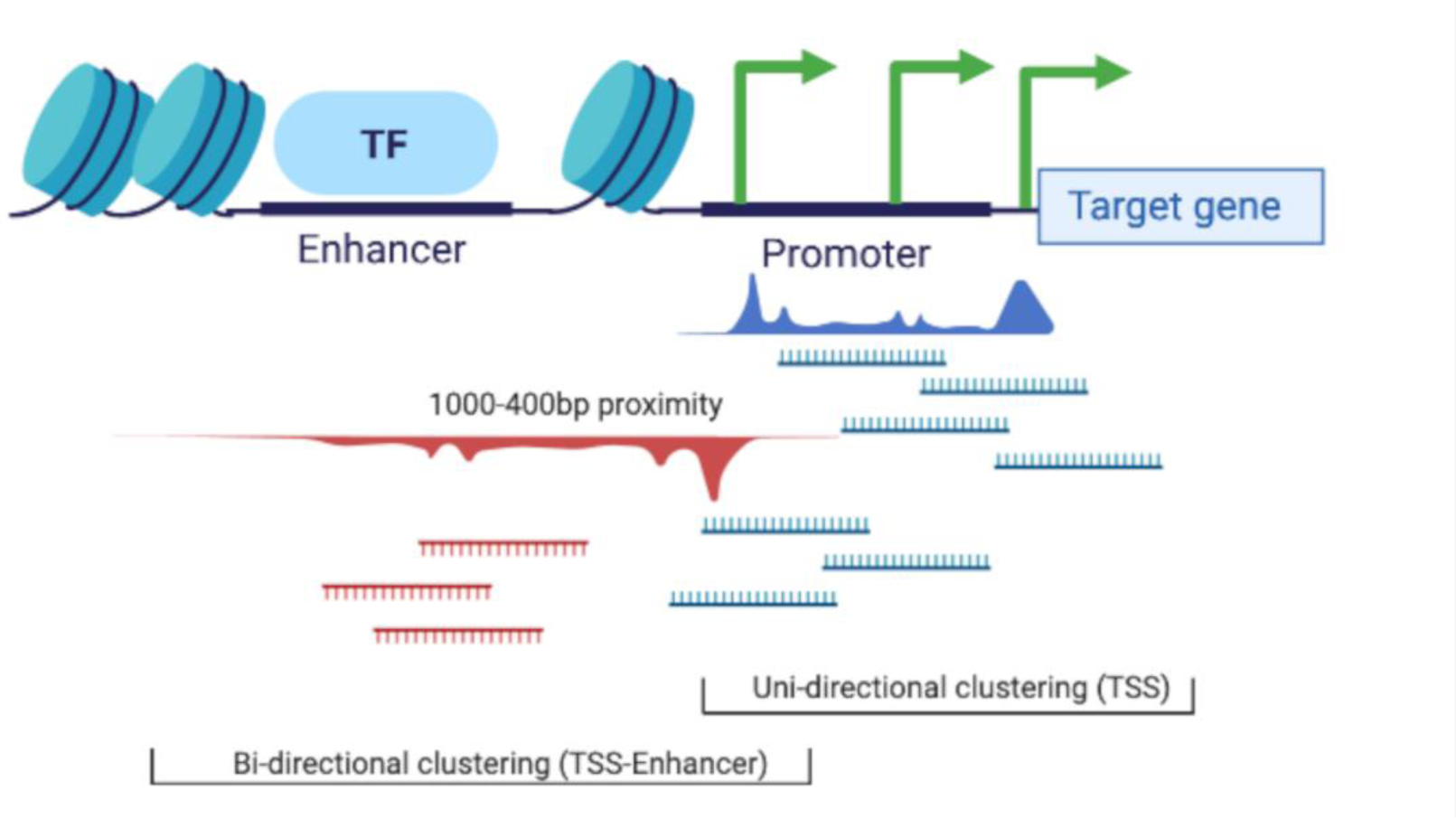
Schematic representation of the two clustering algorithms used in the CAGEfightR package for TSS (uni-directional) and TSS-Enhancer (bi-directional) clustering.

#### Identification of shared TSS or TSS-enhancer clusters across tissues

TSS or TSS-Enhancer clusters that were shared across tissues, were identified by investigating the CTPM expression profile of each of the tissues using correlation based and mutual information (MI) distance matrices (Priness et al., 2007; Reshef et al., 2018). This method of MI based clustering tolerates missingness and outlier-induced grouping errors in gene expression profiles (Priness et al., 2007). Using this method, we assumed that the CTPM expression profile, for each cluster, could vary across tissues. However, for a predicted TSS or TSS-Enhancer cluster to be considered high-confidence and associated with widely expressed genes and widely used promotors, it must be present in at least two thirds of the tissues (37/56) in the dataset.

#### Identification of tissue-specific TSS or TSS-enhancer clusters

The 2/3^rds^ representation threshold applied above would remove all tissue-specific CAGE tag clusters. To overcome this, a rerun of the clustering algorithm was performed with the 2/3^rds^ representation threshold removed. Tissue-specific uni- directional TSS clusters that were only present in 1/56 tissues were identified by filtering for CAGE tags with >10 expressed counts to create a data frame. The data frame was then filtered tissue-by-tissue to only retain uni-directional TSS clusters present in each tissue separately. This process was then repeated for the TSS-Enhancer clusters.

#### Annotation of ‘novel’ TSS in the ovine genome

We expected given the diversity of tissues sampled that we would detect a significant number of ‘novel’, previously unannotated TSS. The CAGE tag uni-directional clusters (TSS) were annotated using the mergeByOverlay function of the GenomicFeatures package in R and the custom TxDB object as following:

*mergeByOverlaps(subject = TSS, query = promoters(txdb, upstream = 25, downstream = 25, use.names = T,c(“tx_name”, “GENEID”)), maxgap = 25, type = “any”).* The TxDB object calculates the range of the promoter based on the 5’UTR and first CDS codon coordinates. In each tissue any putative TSS region within 50bp range of the promoter coordinate of a gene model was considered ‘annotated’. In addition, we expanded this range to 400bp to determine whether this would identify significantly more unannotated TSS further from the promoter. A reverse sub setting of the 50bp window region was performed as follows: *subsetByOverlaps(x = TSS, ranges = annotated, invert = TRUE).* These regions were considered ‘novel’ TSS previously unannotated in the assembly. This process was repeated for every tissue separately (n=56).

### Comparative analysis of WGBS and CAGE Data

#### Preparation of genomic DNA from tissue

Extraction of DNA for bisulphite sequencing was performed using phenol:chloroform:isoamyl alcohol method. Briefly, approximately 1 g frozen tissue was pulverized and resuspended in 2.26ml of digestion buffer (10 mM Tris-HCl, 400 mM NaCl, 2 mM EDTA, pH 8.0) with 200 μl of SDS 10% and 60μl RnaseA (10 mg/ml) (Sigma-Aldrich, St. Louis, MO, USA) RNA degradation proceeded for one hour at 37°C with gentle shaking. Next, 25 μl of proteinase K (20 mg/ml) (Sigma-Aldrich) was added to the suspension and incubated overnight (approximately 16 hours) at 37°C with gentle shaking. The viscous lysate was transferred to a 2ml Phase Lock tube (VWR, Radnor, PA) and extracted twice with Tris-HCl-saturated phenol:chloroform:isoamyl alcohol (25:24:1) pH 8.0, followed by extraction with 2.5ml chloroform. The DNA was precipitated by addition of 5.5 ml of 100% ethanol and 250 μl of 3M Sodium Acetate to the aqueous phase in a 15 ml conical tube, mixed by gentle inversion until the DNA became visible. The DNA was removed with a bent Pastuer pipette hook, washed in 5 ml 70% cold ethanol, air dried then resuspended in 250 μl –1 ml of 1X TE and stored at −20°C until use. DNA concentration was quantified fluorometrically on the Qubit® 3.0 Fluorometer (Thermo Fisher Scientific, Waltham, MA, United States) using the Qubit dsDNA HS Assay Kit. The purity of the extractions was determined via 260/280 and 260/230 ratios measured on the NanoDrop 8000 (Thermo Fisher Scientific) and DNA integrity was assessed by 1% agarose gel electrophoresis. The protocol is available via the FAANG Data Coordination Centre https://data.faang.org/api/fire_api/assays/USDA_SOP_DNA_Extraction_From_WholeBloodandLiver_20200611.pdf.

#### Whole Genome Bisulphite Conversion and Sequencing

Library preparation and sequencing of seven tissues and 1 cell type (Figure 1), selected to include a representative from all major organ systems, were performed by The Garvan Institute of Medical Research, Darlinghurst, Sydney, New South Wales. Un-methylated lambda DNA was added at 0.5% of the total sample DNA concentration prior to bisulphite conversion as a conversion efficiency control. DNA conversion was carried out using the EZ DNA Methylation-Gold Kit (Zymo Research, CA, USA) following the manufacturer’s instructions. The Accel-NGS Methyl-seq DNA kit (Swift Biosciences, MI, USA) for single indexing, was used to prepare the libraries, following the manufacturer’s instructions. Libraries were pooled together and sequenced across 6 lanes of a flow-cell on an Illumina HiSeq X platform using paired end chemistry for 150 bp reads (min 10X coverage). The protocol is available via FAANG Data Coordination Centre https://data.faang.org/api/fire_api/assays/AGR_SOP_WGBS_AgR_Libary_prep_20200610.pdf

#### WGBS data processing

Paired end Illumina WGBS sequence data were processed and analysed using in house scripting (bash and R) and a range of purpose-built bioinformatics tools on the AgResearch and University of Edinburgh high performance computing facilities. The analysis protocol for WGBS is available via the FAANG Data Coordination Centre https://data.faang.org/api/fire_api/analysis/AGR_SOP_WGBS_AgR_data_an <u>alysis_20200610.pdf</u> and summarised below.

Briefly, FASTQ files for each sample, run across multiple lanes were merged together. TrimGalore v. 0.5.0. (https://github.com/FelixKrueger/TrimGalore) was used to trim raw reads to remove adapter oligos, poor quality bases (phred score less than 20) and the low complexity sequence tag introduced during Accel-NGS Methyl-seq DNA kit library preparation as follows: *trim_galore -q 20 --fastqc --paired --clip_R2 18 --three_prime_clip_R1 18 --retain_unpaired –o Trim_out INPUT_R1.fq.gz INPUT_R2.fq.gz* A bisulphite-sequencing amenable reference genome was built using the *Oar rambouillet v1.0*, GenBank Accession number: GCA_002742125.1 genome with the BSSeeker2 script *bs_seeker2-build.py* using bowtie v2.3.4.3 (Langmead and Salzberg, 2012). and default parameters. The Enterobacteria phage lambda genome available from NCBI (Accession number: NC_001416) was added to the Oar rambouillet *v1.0* genome as an extra chromosome to enable alignment of the unmethylated lambda DNA conversion control reads. Paired-end, trimmed reads were aligned to the reference genome using the BSSeeker2 script *bs_seeker2-align.py* and bowtie v2.3.4.3 (Langmead and Salzberg, 2012) allowing four mismatches (-m 4). Aligned bam files were sorted with samtools v1.6 (Li et al., 2009) and duplicate reads were removed with picard tools v2.17.11 (https://broadinstitute.github.io/picard/) MarkDuplicates function.

Deduplicated bam files were used to call DNA methylation levels using the “bam2cgmap” function within CGmaptools (Guo et al., 2018) with default options to generate ATCGmap and CGmap files for each sample. The ATCGmap file format summarises mapping information for all covered nucleotides on both strands, and is specifically designed for BS-seq data; whilst the CGmap format is a more condensed summary providing sequence context and estimated methylation levels at any covered cytosine in the reference genome.

Hyper-methylated and hypo-methylated regions were determined for each sample using methpipe v3.4.3 (Song et al., 2013). Specifically, CGmap files for each sample were reformatted for the methpipe v3.4.3 workflow using custom awk scripts. The methpipe symmetric-cpgs program was used to merge individual methylation levels at symmetric CpG pairs. Hypo-methylated and hyper-methylated regions were determined using the hmr program within methpipe, which uses a hidden Markov model (HMM) using a Beta-Binomial distribution to describe methylation levels at individual CpG sites, accounting for the read coverage at each site.

Visualisation of the individual CpG site methylation levels with a minimum read depth cut-off of 10x coverage was done using Gviz package v.1.28.3 (Hahne and Ivanek, 2016).

#### Comparative analysis of annotated and ‘novel’ TSS with WGBS methylation information

We expected that reproducible TSS, either annotated or novel, would overlap with hypo-methylated regions of the genome (Yamashita et al., 2005; Yagi et al., 2008). To test whether this was true for those identified in our analysis, both annotated and novel TSS from the CAGE BED tracks were intersected with WGBS hypo methylation profiles using bedtools v.2.29.2 (Quinlan and Hall, 2010) and the following script: *bedtools intersect -b WGBS_HypoCpG.bed -a Novel_or_ Annotated.bed > Novel_or_annotated_HypoCpG.bed.* Any annotated and novel TSS (within a ±50bp window of the promoter) that intersected hypomethylated regions of DNA in each tissue, were verified as reproducible TSS and the remainder as ‘noise’. The overlay of these regions was visualised as a genomic track using the Gviz package v.1.28.3 (Hahne and Ivanek, 2016).

#### Visualisation of the annotated TSS, mRNA-Seq and WGBS tracks in the ovine genome

In order to confirm the simultaneous expression of mRNA, CAGE tags corresponding to an active TSS and a hypomethylated region of DNA, a genomic track on which all three datasets could be visualised was generated. This visualisation consists of the following tracks: 1) Uni-directional CAGE tag clusters (TSS) 2) Bi-directional CAGE tag clusters (TSS-Enhancers), 3) WGBS hypomethylation score (bp resolution), 4) Transcript level expression (mRNA-Seq [TPM]), 5) The transcript models, and 6) The gene model. Areas of the genome where TSS or TSS-Enhancer regions overlapped regions with a high hypomethylation score, within 5’ end of an actively expressing transcript (TPM score), were considered reproducible TSS for that tissue. This process was performed using eight tissues with matching mRNA-Seq, CAGE and WGBS sequence data. The Gviz package v.1.28.3 was used to visualise these tracks (Hahne and Ivanek, 2016).

#### Validation of tissue-specific expression profiles mRNA-Sequencing

Total RNA for mRNA-Seq from 32 tissues (Figure 1) was prepared, as above for the CAGE samples, by USMARC, and for 26 tissues by Baylor College of Medicine (BCM) using the MagMAX mirVana total RNA isolation kit (Thermo Fisher Scientific, Waltham, MA, United States) according to the manufacturer’s instructions. Paired end polyA selected mRNA-Seq libraries were prepared and sequenced on an Illumina NextSeq500 at USMARC or the Ilumina HiSeq2000 at BCM using the Illumina Tru- Seq Stranded mRNA Library Preparation Kit. For each tissue a set of expression estimates, as transcripts per million (TPM), were obtained using the transcript quantification tool Kallisto v0.43.0 (Bray et al., 2016). The mRNA-Seq analysis pipeline is accessible via the FAANG Data Coordination Centre https://data.faang.org/api/fire_api/analysis/ROSLIN_SOP_RNA-Seq_analysis_pipeline_20200610.pdf. A pairwise distance matrix (multiple correlation coefficient based) was produced using MI values for all tissues and a dendrogram of tissues was created in order to visualise grouping patterns of tissues with similar mRNA expression profiles, and for comparison with the CAGE dataset.

#### Comparative analysis of tissue-specific expression profiles using information from CAGE and mRNA-Seq

We assessed whether TSS expression profiles from the CAGE dataset were biologically meaningful using the mutual information (MI) sharing algorithm (Joe, 1989). Tissues with the same function and physiology should have similar TSS expression profiles. The CTPM expression level was binned (n=10) using the bioDist package v.1.56.0 (Ding et al., 2012) and mutual information (MI) for each pair of tissue samples was calculated as in (Joe 1989). :

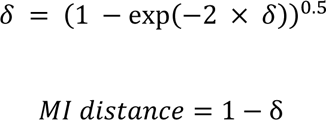

A pairwise distance matrix (multiple correlation coefficient based) was produced using MI values for all tissues and a dendrogram of tissues created to visualise grouping patterns of tissues with similar TSS expression profiles. If the expression profiles were meaningful then tissues with similar function and physiology would group together in clades within the dendrogram. These tissue specific groupings were then further validated by comparison with mRNA-Seq data for the same samples, using the MI sharing algorithm and dendrogram approach.

## Results

### Library size and annotation metrics

The mean CAGE library depth based on uniquely mapped CAGE reads was 4,862,957 reads. A detailed explanation of the attrition of reads at each stage of the analysis pipeline is included in Supplementary File 1, Section 1.1. Library depth varied across tissues. Tissues with low depth were not related to any specific barcodes and were evenly spread over the two sequencing runs (Supplemental Table S1, Supplementary Figure S1), suggesting random variation rather than systematic differences due to specific barcodes or sequencing runs. The RIN^e^ values were also consistently >7 for all tissues with low counts, indicating RNA integrity was also unlikely to be affecting library depth. Differences in tag numbers are therefore more likely to relate to variation in efficiency between individual libraries or tissue-specific differences related to the physiology of the tissue.

### CAGE tag clustering and annotation by genomic regions

We used a newly developed software package to annotate TSS in the Rambouillet Benz2616 genome (Thodberg and Sandelin, 2019; Thodberg et al., 2019) which clustered the CAGE tags as A) uni-directionally into predicted TSS or B) bi-directionally into correlated TSS and enhancer (TSS-Enhancer) clusters (Figure 3). The clustered CAGE tags were filtered to remove any clusters with a minimum expression level of <10 tag counts. The mean (±SD) and median number of tissues per cluster was 3.68 ± 4.78 and 2, respectively. Application of the 2/3^rds^ representation criteria (i.e. a minimum of 37/56 tissues had to express the tag cluster) and filtering out of tag clusters with <10 TPM resulted in an average of 8,219 uni-directional TSS clusters, from a total of 5,450,864 (pre-filtering), for downstream analysis. A detailed description of the cluster metrics at each stage of filtering is included in Supplementary File 1 Sections 1.1. and 1.3. Although direct comparisons are difficult due to differences in methodology and the relative ‘completeness’ of the reference annotation used the level of retained CAGE sequencing datasets (0.5% retained clusters with 2/3^rds^ tissue representation) is somewhat lower than reported for other mammalian promoter-level expression atlas projects. In the FANTOM5 project, for example, approximately 5% of clusters were retained (Forrest et al., 2014a). To further validate the 2/3^rds^ tissue representation criteria we chose we also investigated the number of transcripts captured in the poly-A enriched mRNA-Seq dataset. Poly-A enriched mRNA-Seq data was available for 52 matching tissues and captured a smaller number of transcripts (n=32,852) with TPM>10 in comparison to CAGE CTPM>10 (n=53,507). Direct comparison of expression for CAGE tags (27nt) and paired end RNA-Seq (75nt) reads could result in technology dependent bias. Taking this into consideration the CAGE dataset with the 2/3^rds^ representation criteria applied provided annotation for 31,113 transcripts with minimum CPTM>10. When the same criteria was applied to the mRNA-Seq dataset only 3,908 transcripts with TPM>10 were annotated. The expression (as TPM) of transcripts for each tissue and the TPM threshold metrics are included in Supplementary file1 and Supplementary Table 4.

Bi-directional TSS-enhancer clusters were far fewer in number, although retention was higher with over 23% meeting the same 2/3^rds^ representation criteria 741 from a total of 3,131. Though fewer in number these bi-directional (including TSS-enhancer) clusters are functionally important in the regulation of expression of their target genes (Andersson et al., 2014; Thodberg and Sandelin, 2019), consistent with finding them in over 2/3^rds^ of tissues. The co-expression of leading enhancer RNA (eRNA) which is captured by CAGE sequencing can provide a map to enhancer families in the genome and the genes under their regulation (Andersson et al., 2014).

The locations of both uni-directional TSS and bi-directional TSS-enhancer clusters were identified in *Oar rambouillet v1.0* and the proportion of TSS clusters located within or near annotated gene features was estimated (Figure 4). The custom BSgenome and TxDB objects created from the GFF3 file format provide detailed calculated coordinates for the following sections: intergenic (>1000bp before 5’UTR or after the end of 3’UTR), proximal (1000bp upstream of the 5’UTR), promoter (±100bp from 5’UTR) and the standard gene model (5’UTR, exon, intron and 3’UTR). The genomic region class with the highest number of unidirectional clusters (39.25%) was the promoter regions (±100bp from 5’UTR) (Figure 4A), with a relatively even distribution within the other regions of the genome, including 6% mapping proximally to the 5’UTR. The majority of bi-directional TSS-enhancer clusters were also located in promoter regions (70.1%) with a smaller proportion (25.6%) located in proximal regions (Figure 4B). The lack of bi-directional TSS-enhancer clusters in other regions is a consequence of the operation of the CAGEfightR algorithm, which only considers bi-directional clusters within a 400-1000bp window of a TSS CAGE tag cluster (Thodberg and Sandelin, 2019; Thodberg et al., 2019). This approach also reduced the total count compared to unidirectional clusters (28,148 uni-directional clusters relative to 741 bidirectional TSS-enhancer clusters across tissues) (Thodberg et al.,2019).

**Figure 4.**
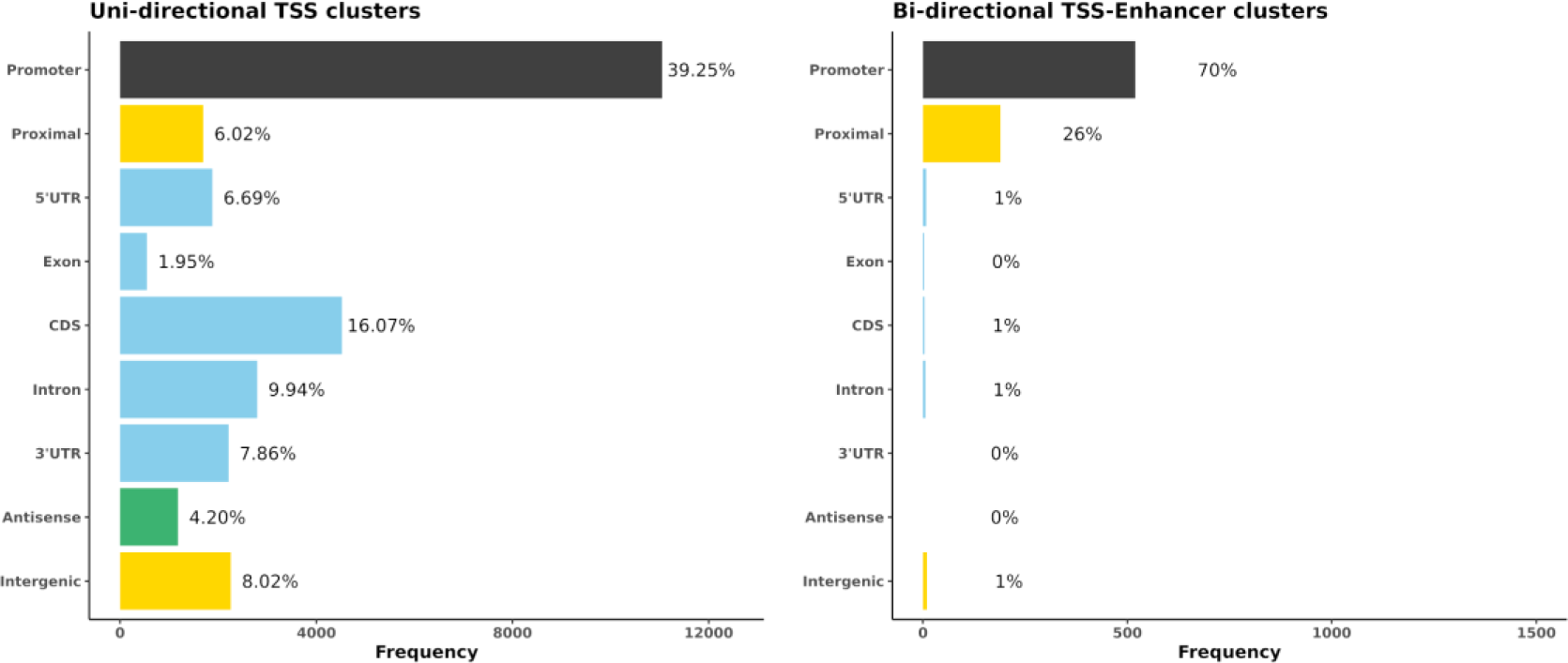
The genomic region distribution of CAGE tag clusters mapped against *Oar rambouillet v1.0* assembly and gene annotation. The counts were averaged across tissues. A) Uni-directional TSS clusters with the highest proportion in promoter region (±100bp of the 5’UTR beginning at the [TSS]). B) Bi-directional TSS-enhancer clusters with the highest proportion in the proximal region (1000bp upstream of the 5’UTR beginning at the [TSS]).

### Capturing metrics of CAGE tag clusters per gene

During the clustering process we also determined the proportion of annotated genes and transcripts in the Oar rambouillet v1.0 NCBI annotation that we did not capture using our dataset. When the 2/3^rds^ representation filtering criteria was applied 44.7% of transcripts (25,195) and 54.6% of genes (16,950) were not captured by our CAGE TSS clusters. When the 2/3^rds^ representation filtering criteria was removed, and presence of the CAGE tag in only one tissue out of 56 considered sufficient, the proportion that we didn’t capture was reduced to 7% of genes and 5% of transcripts.

To investigate whether some genes posessed multiple putative TSS we also estimated the number of CAGE TSS clusters per gene. The median and also the highest frequency of TSS cluster per gene was 1 (mean 1.8) (13,912 genes / 31,3113 transcripts annotated using our dataset), indicating that the vast majority of genes annotated in the Oar rambouillet v1.0 reference genome have only one TSS, and genes with more than five TSS were rare (Supplementary figure S5 and Supplementary Table 5).

### Distribution of CAGE tag clusters in *Oar rambouillet v1.0* relative to *Oar_v3.1*

The new reference sheep genome assembly (Oar rambouillet v1.0) is more contiguous than the earlier draft genome sequence Oar_v3.1 (Jiang et al., 2014) with contig N50 values of 2,572,683 bp and 40,376 bp, respectively and would be expected to provide a better template for annotation of gene models and other genomic features. As a proxy for testing this assumption we investigated how mapped CAGE tag clusters were distributed across the two genome assemblies (Supplemental Figure S2). The percentage of uni-directional CAGE tag clusters mapping to intergenic regions, which usually occurs due to missing gene model information, was greater for *Oar_v3.1* (33.9%) relative to *Oar rambouillet v1.0* (8%). The percentage of uni-directional CAGE tag clusters mapping to annotated promoter regions was greater for *Oar rambouillet v1.0* (39.25%) compared to *Oar_v3.1* (14.94%), indicating the proportion of accurate gene models in *Oar rambouillet v1.0* was greater. Of the 28,148 unidirectional TSS clusters mapped to *Oar rambouillet v1.0,* 87.74% mapped to 13,868 unique genes (31,729 transcripts). In comparison, of the 23,829 unidirectional TSS clusters mapped to *Oar_v3.1,* 49.1% mapped to 6,549 genes (9,914 transcripts). A larger number of TSS-Enhancer CAGE clusters were detected in *Oar_v3.1* (1121) in comparison to *Oar rambouillet v1.0* (741) mapping to 1371 and 2598 unique genes, respectively. A detailed comparison of mapping of the CAGE tags to the two reference assemblies is included in Supplementary File 1, Sections 2 and 3.

### Mapping of CAGE tags shared across all tissue samples

Correlation-based and mutual information (MI) distance matrices were used to evaluate the occurrence of TSS and enhancer TSS across tissues. The mean ± SD number of tissues in which each cluster passing the 2/3^rds^ criteria (expressed in 37/56 tissues) was (47.73 ± 6.03). Uni-directional TSS clusters (n=28,148 TSS regions) that were shared across tissues and detected in at least 37/56 tissues are visualised in Figure 5. Each chord in Figure 5 represents the presence of an expressed unidirectional TSS cluster shared across tissues. The majority of the unidirectional TSS that were shared across tissues mapped to promoters (39.25%) and were shared evenly across the tissues sampled (Figure 5). Some tissues e.g. mammary gland, pituitary gland and urinary bladder had more uni-directional TSS mapping to intergenic regions, which might indicate evidence of alternative splicing or differential TSS usage across tissues (Figure 5). Alternative splicing events and differential TSS usage, captured by CAGE, are often not included in the reference gene prediction models (Berger et al., 2019).

**Figure 5.**
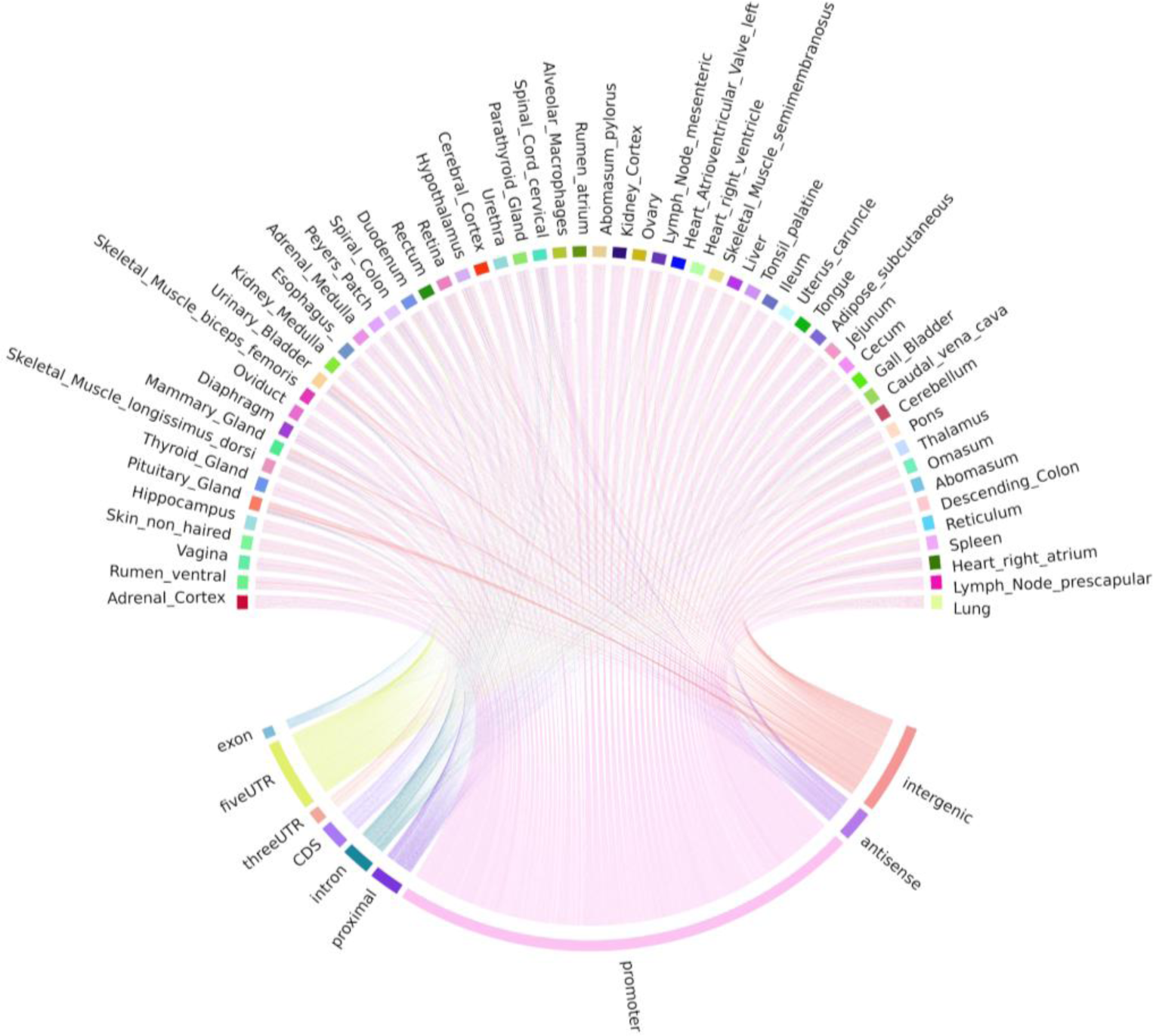
Chord diagram of expression level (TPM) of CAGE tag clusters (uni- directional TSS) across all the tissues collected from Benz2616. Shared CAGE tag clusters are common to at least 2/3^rds^ of the tissues (37/56).

Bi-directional TSS-Enhancer CAGE clusters were far fewer in number but were shared in a similar pattern across tissues as the uni-directional TSS clusters (Figure 6). The majority (70.1%) of the TSS-Enhancer clusters mapped to promoters (n=520) while 25.6% mapped to ‘proximal’ regions as expected according to the 400bp-1000bp detection window for TSS-Enhancer clusters from the centre of the promoter (Figure 6). For some tissues including abomasum, spleen and heart right atrium the proportion of bi-directional TSS-Enhancer clusters mapping to proximal regions was greater indicating more enhancer families could be present within these tissues (Figure 6).

**Figure 6.**
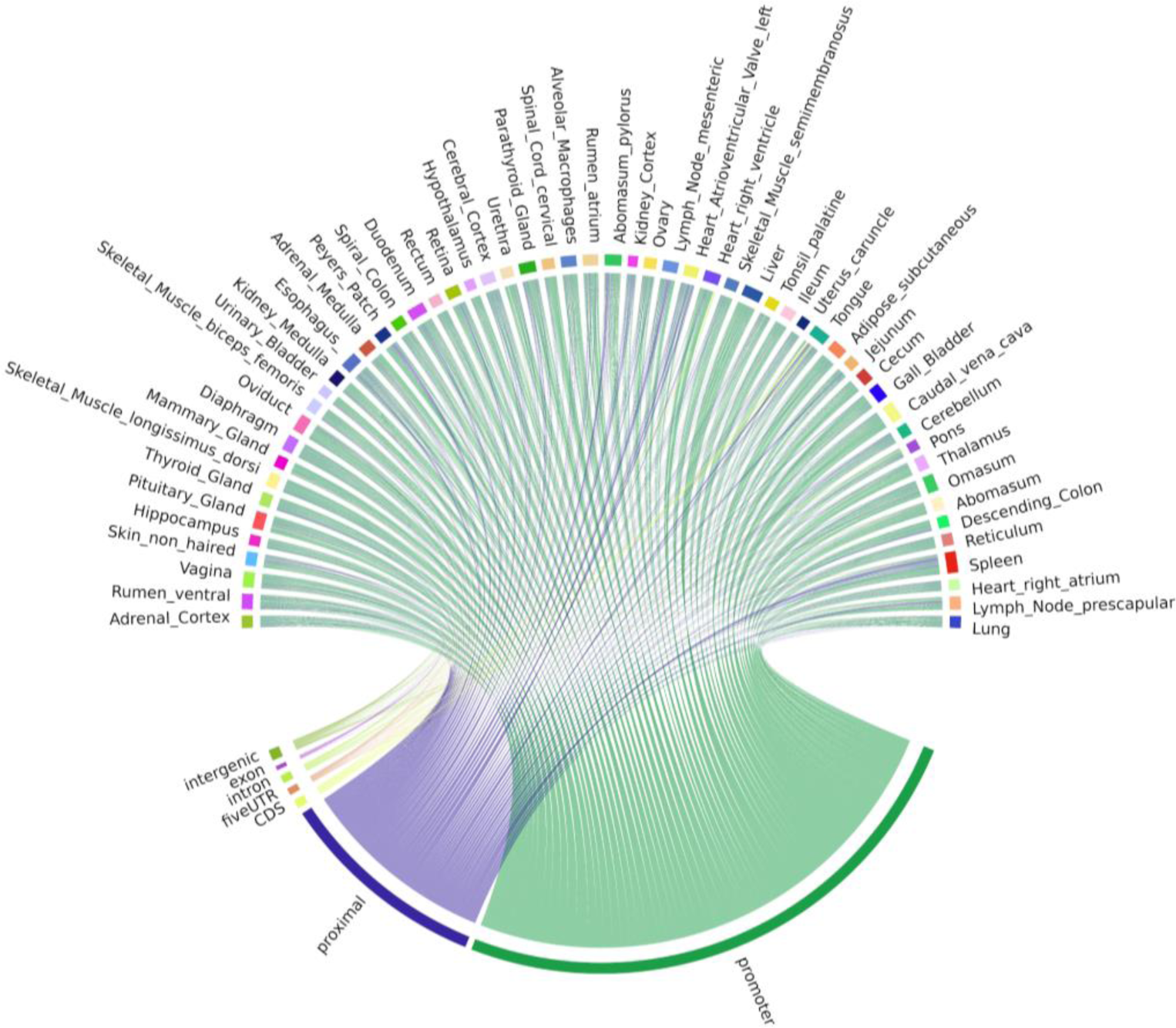
Chord diagram of expression level (TPM) of CAGE tag clusters (bi- directional TSS-Enhancer) across all the tissues collected from Benz2616. CAGE tag clusters expressed (>10 CTPM) by at least 2/3^rds^ of the tissues (37/56).

### Mapping of tissue-specific CAGE tags

The application of the 2/3^rds^ criteria provided a high level of confidence in assigning TSS and TSS enhancer elements, but eliminated the ability to observe potential tissue-specific CAGE tags or TSS clusters. Tissue-specific tags, i.e. those observed in only one of the 56 tissues, were examined to evaluate the ability to distinguish tissue-specific clusters from the background. A total of 3,228,425 tags were observed in only one tissue, and a much higher proportion (80.0%) of these tags mapped to intergenic and intronic regions compared to tags found across tissues, suggesting they do not represent true TSS (Supplementary Table 2). Only 0.8% of the tissue-specific CAGE tag clusters mapped to promoter or proximal regions (Supplementary Table 2). The caecum (n=1554), cerebellum (n=601) and longisimus dorsi muscle (n=477) had the highest number of tissues-specific predicted unidirectional TSS. The greatest number of expressed TSS (>1 CTPM) were detected in ceberellum (84/601) as shown in Supplementary Figure S3A. However, the expression level of tissue-specific CAGE tag clusters was very low (<2 CTPM), which combined with the small sample size (n=1) for each tissue, meant that analysis of tissue-specific TSS was not particularly meaningful using this dataset. The analysis was repeated for tissue specific TSS-Enhancer clusters which is detailed in Supplementary Figure S3B.

### Proportion of ‘novel’ TSS within the CAGE dataset for each tissue

CAGE tag clusters were annotated initially using the *Oar rambouillet v1.0* gene models from NCBI. A tissue-by-tissue annotation was performed using the same gene models to identify any CAGE tag clusters within a 50bp window of the promoter boundaries of every gene. From a total of 23,994 ± 518 TSS (the average number of TSS per tissue ± SE) we found 11,349 ± 170 (49.8% ± 0.01) were located within 50bp of the promoter. The CAGE tag clusters were annotated using the NCBI *Oar rambouillet v1.0* GFF3 gene track file (version 103) and a TxDB object created in the GenomicFeatures package (version 1.36.4) in R. CAGE tag clusters within 50bp (short range) or 400bp (long range) of the promoter were defined as annotated. Supplementary File 2 includes BED files for these CAGE tag clusters. The percentage of ‘novel’ previously un-annotated, but likely to be reproducible, CAGE tag clusters for each tissue within 50bp (short range) and 400bp (long range) from the promoter are detailed in Table 1.

**Table 1:**
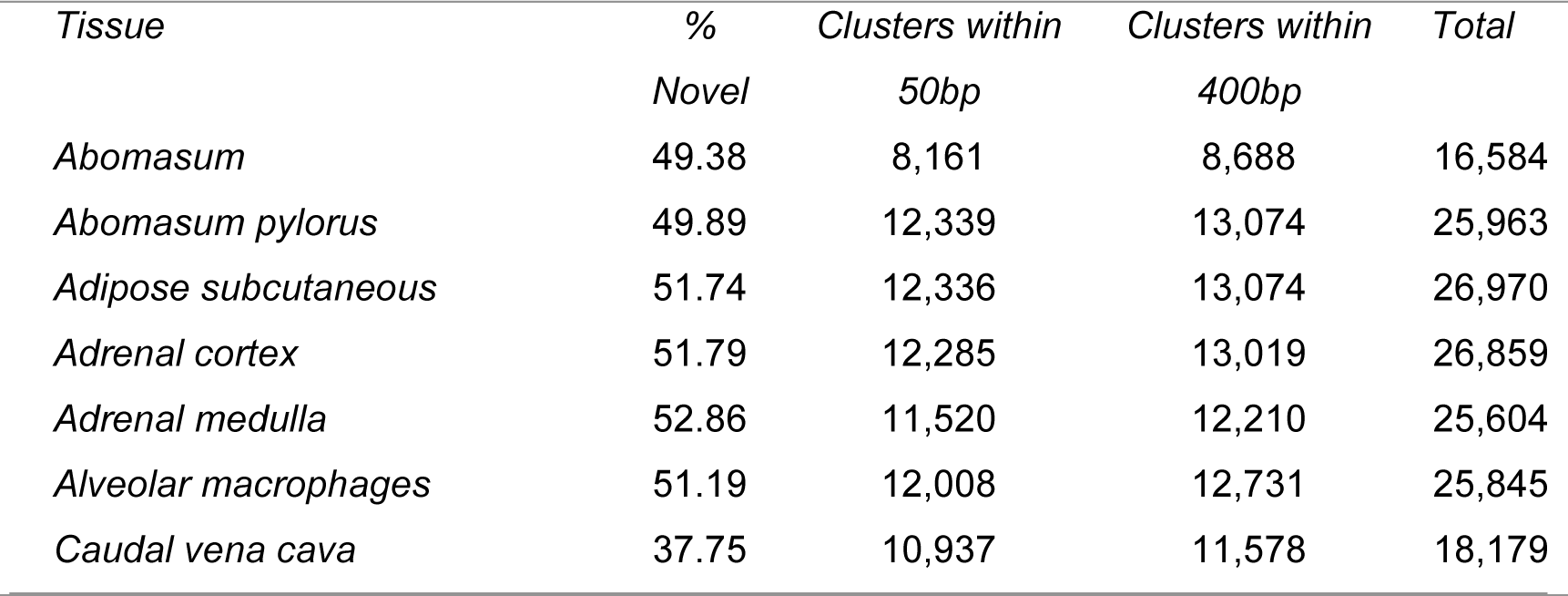
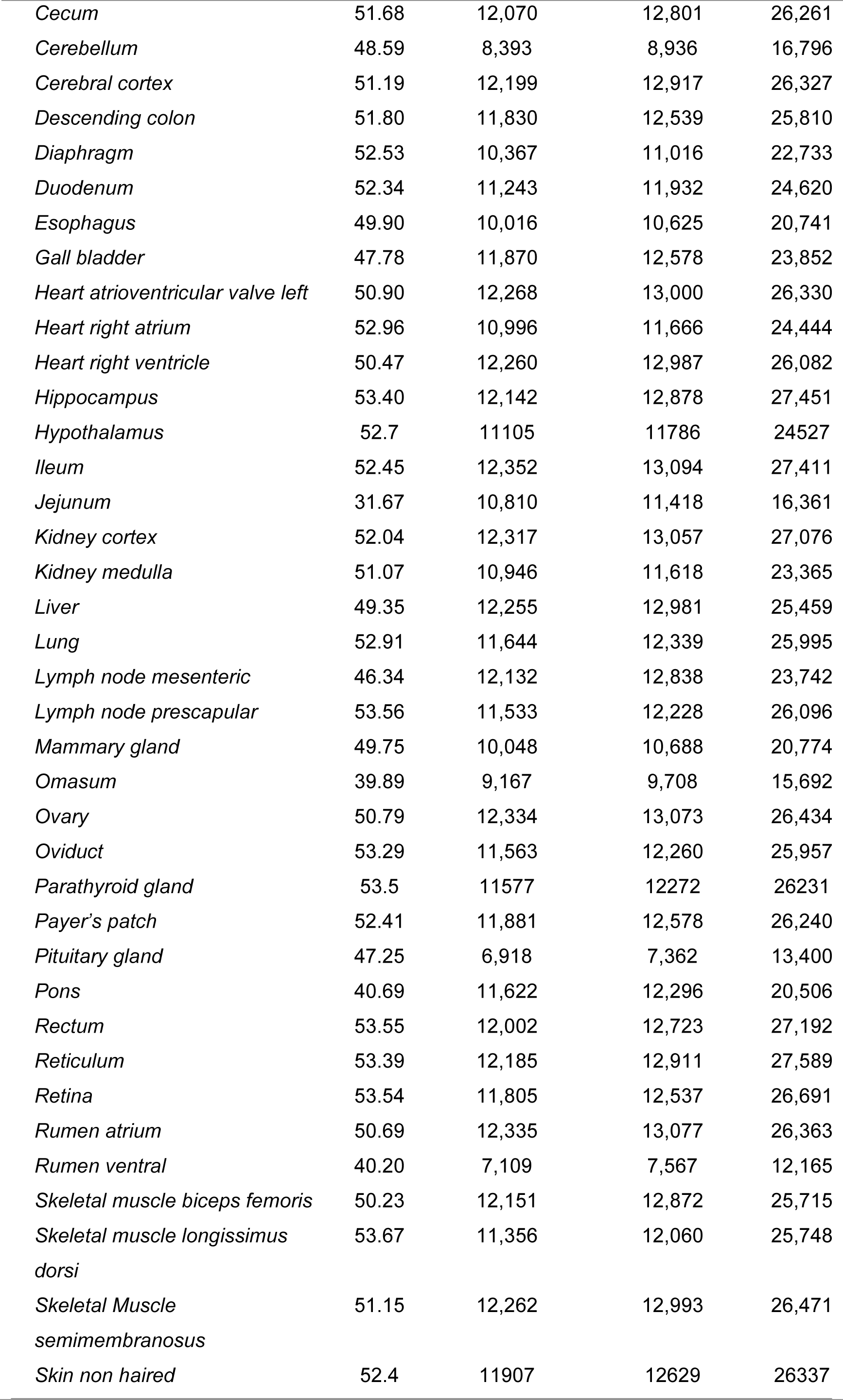
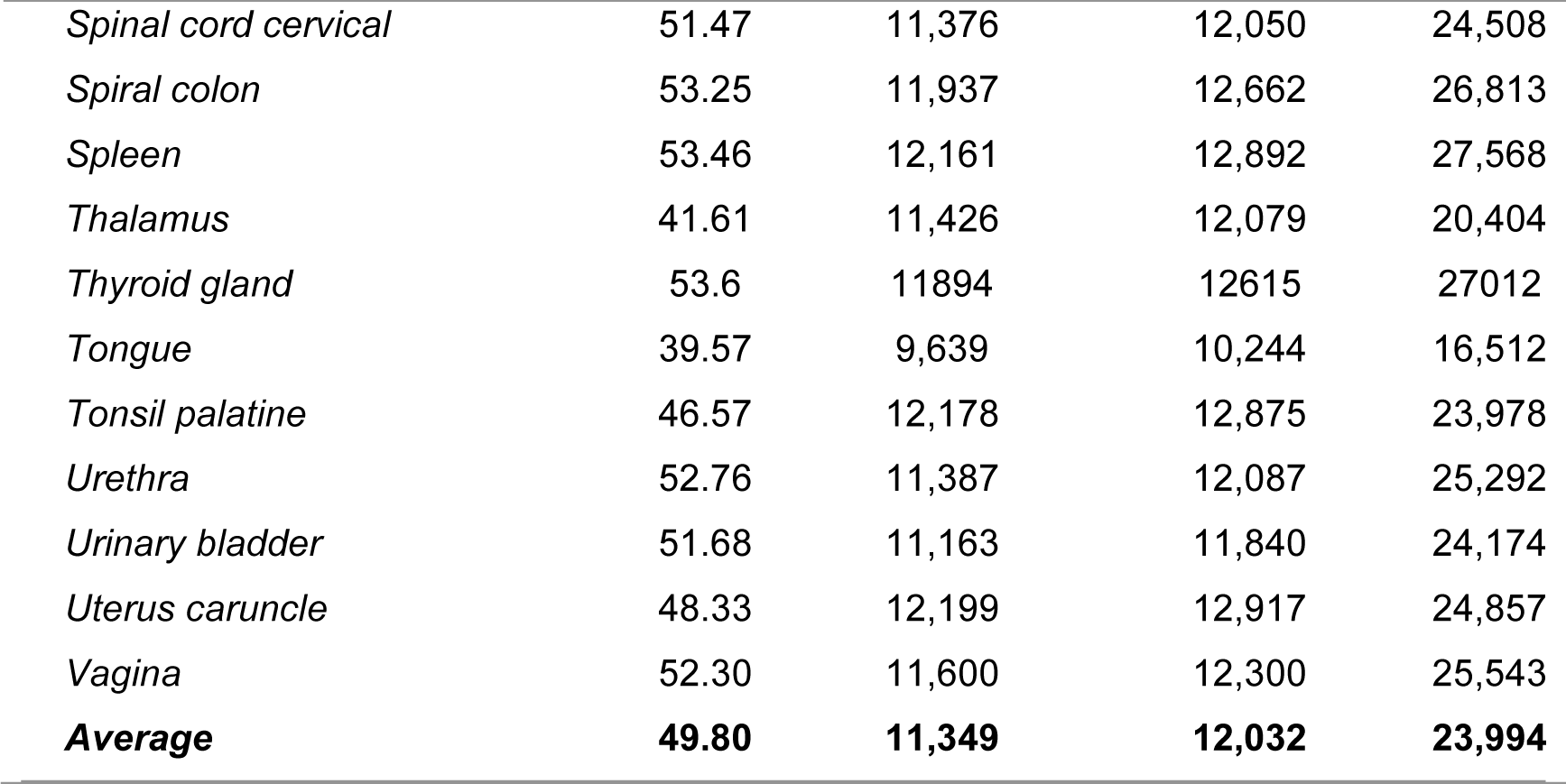
The total number and percentage of ‘novel’ CAGE tag clusters for each tissue within 50bp (short range) and 400bp (long range) from the promoter. Total number represents the number of CAGE tag clusters (from the total of 28,148) that are present in each of the 56 tissues. The clusters for each tissue were then annotated by proximity to promoter regions either 50bp or 400 bp (the later includes the count of the former). The % Novel represents the count of the clusters falling outside of 400bp vicinity of any current promoter region.

### Comparative analysis of CAGE and WGBS to validate ’novel’ TSS

True TSS and TSS enhancer elements are very likely to be associated with areas of hypomethylation (Yamashita et al., 2005, Yagi et al, 2008). The assessment of hypomethylation of regions where “novel” TSS were identified thus provides a means to support or question their designation as true TSS. The methylation status of putative TSS regions for eight of the tissues used for CAGE analysis was examined at single nucleotide resolution using WGBS. Each WGBS library was pooled prior to sequencing and multiplexed across eight lanes of the HiSeq X 10 platform. Following trimming of the raw reads, the sequenced libraries produced an average of 103 Gbp of clean data. The average mapping rate of the reads was 78.8%. A small proportion (8.5%) of mapped reads were identified as PCR or optical duplicates and were removed prior to downstream analysis. The average read depth of the filtered libraries was 20x coverage (Supplementary Table S3). Only cytosines with a minimum of ten reads were retained for the subsequent comparative analysis with CAGE data to ensure a high level of confidence in the methylation level estimates, as per published recommendations (Doherty and Couldrey, 2014; Ziller et al., 2015). We would expect that reproducible TSS, either annotated or novel, would overlap with hypo-methylated regions of the genome (Yamashita et al., 2005; Yagi et al., 2008). Comparative analysis of the CAGE data with the WGBS methylation levels from eight tissues from Benz2616 was used to investigate methylation levels at the TSS in comparison to gene body and UTR regions. For the majority of genes, the methylation level was much lower around the transcriptionally active TSS or regulatory enhancer candidate regions compared to the gene body (e.g. for gene *IRF2BP2* Figure 7). We overlaid the WGBS hypomethylated regions and the CAGE uni-directional TSS clusters (annotated and ‘novel’) within 50bp of the promoter. For the eight matching tissues 88.7% of the annotated TSS clusters and 32.2% of the ‘novel’ TSS were hypomethylated (Figure 8).

**Figure 7.**
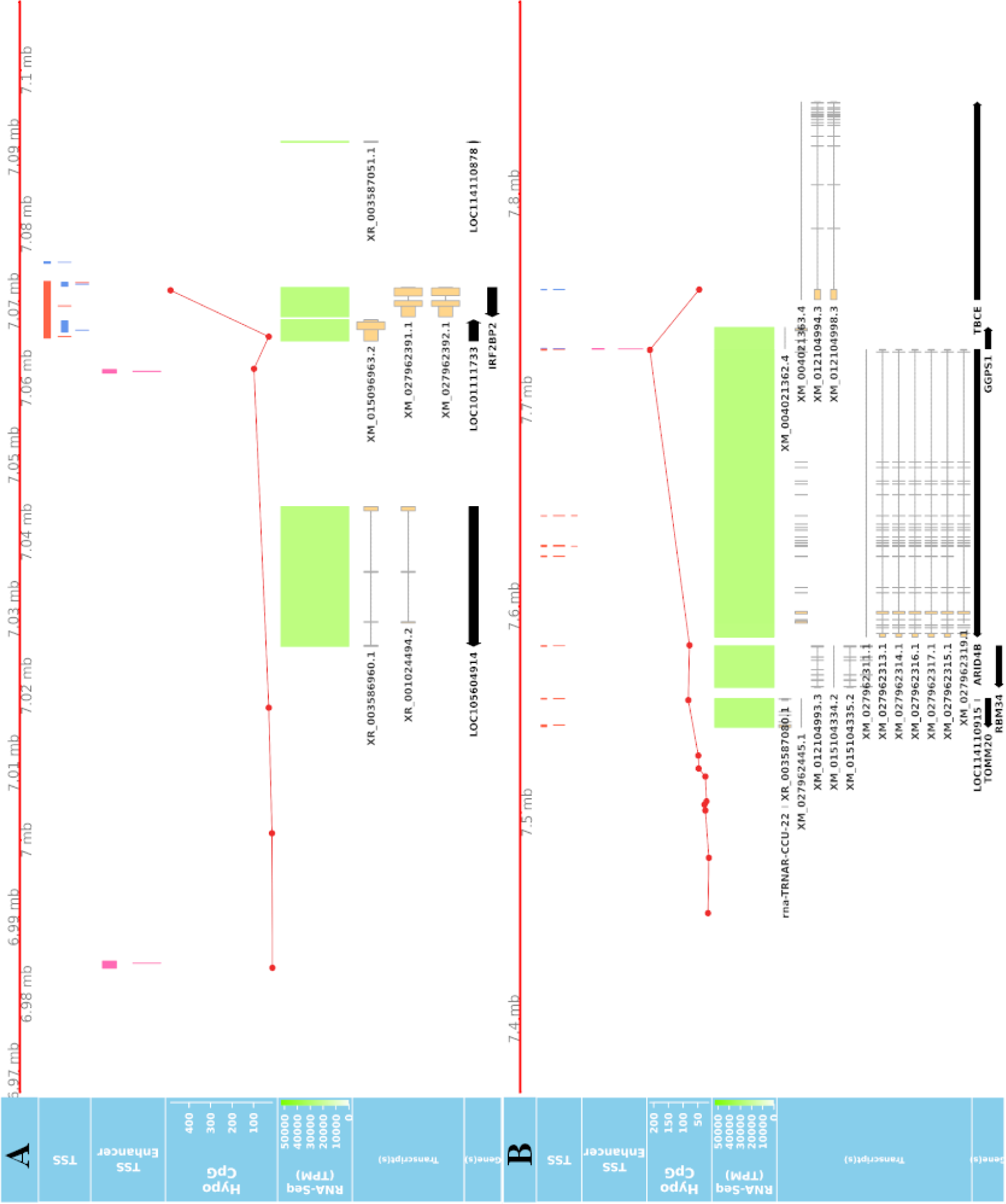
Overlay of CAGE, RNA-Seq and WGBS data tracks centred using the genomic coordinates of genes *IRF2BP2 and ARID4B*. A) Shows a hypomethylated area overlapping multiple uni and bi-directional CAGE tag clusters at 5’UTR of IRF2BP2. B) Predicted CAGE tag clusters with no verifying hypomethylation island within the middle of ARID4B gene, which are likely to be ‘noise’.

**Figure 8:**
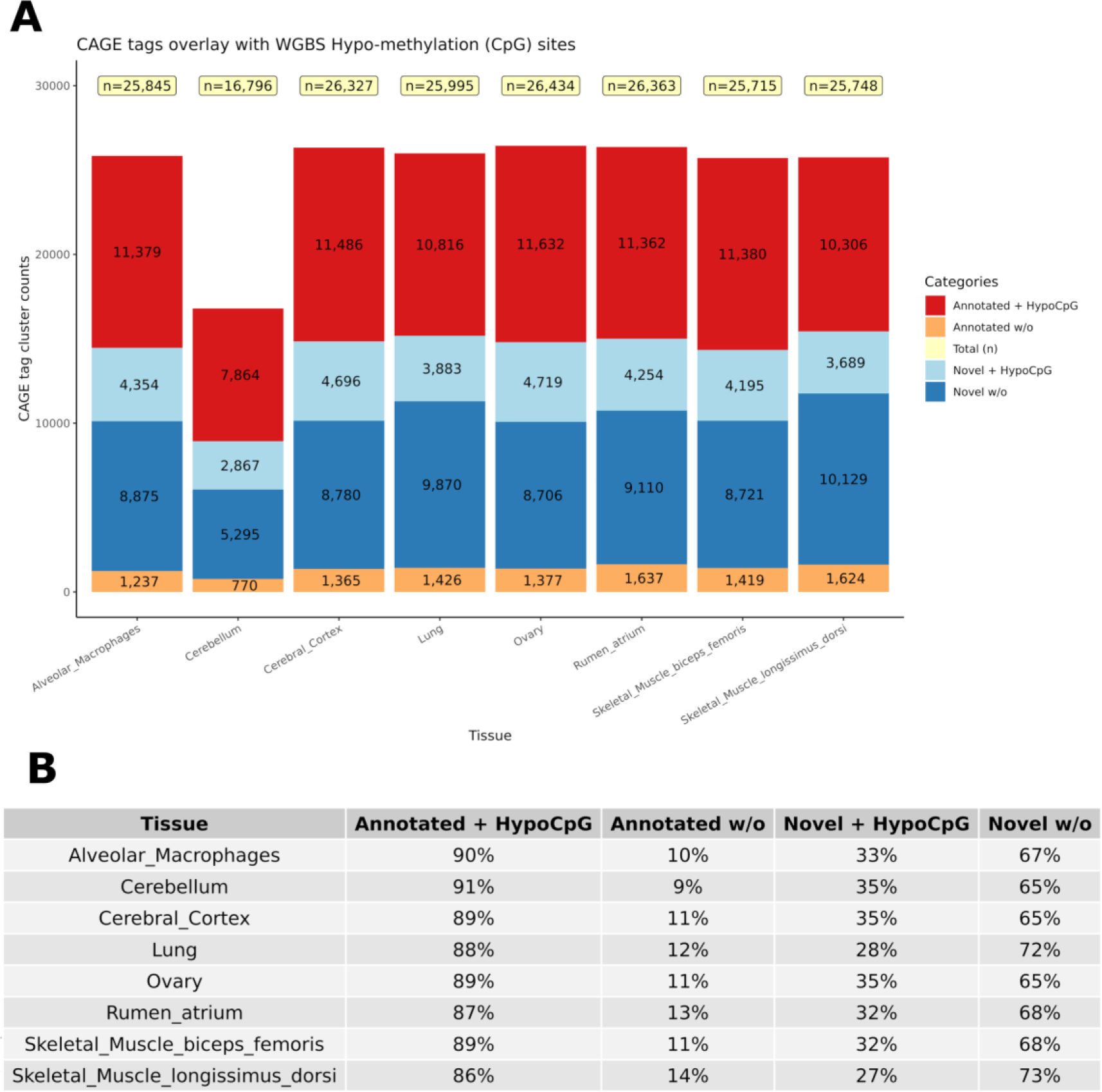
Numbers of CAGE TSS that were hypomethylated according to the WGBS data to distinguish between ‘novel’ reproducible (+HypoCpG) TSS and ‘noise’ (w/o). A) Shows the distribution of CAGE clusters as novel and annotated with or without HypoCpG. B) Percentage of CAGE clusters in each categories for each of the eight tissues.

The combined evidence of the hypomethylation and TSS support the conclusion that 32.2% are in fact novel TSS clusters, whereas 67.8% of the novel TSS clusters lack this confirmation.

### Validation of tissue expression profiles using mRNA-Seq

The tissue samples from Benz2616 were collected for the purpose of annotating her genome and as such N=1 in all cases. As an alternative strategy to having multiple biological replicates we validated the expression profiles for each tissue by comparing the CAGE data (CTPM) and mRNA-Seq (TPM) in 52 matching tissues. The transcript expression TPM was significantly correlated with the CAGE tag cluster CTPM values (correlation coefficient 0.19, Pearson *p* value < 1.0e-08) and visualised as a heatmap (Supplementary Figure S4).

The similarity of tissue expression profiles for the uni-directional TSS clusters was estimated in order to determine if tissues with similar physiology and function formed distinct groups as expected. Similarity (distance) analysis showed a partial grouping based on tissue type and organ system as shown in Figure 9A. Physiologically similar tissues including nervous system and muscle tissues grouped closely together. This grouping was also present in the mRNA-Seq data from tissue matched samples Figure 9B, indicating good correlation between the two datasets. Similar groupings based on organ system and tissue type were observed for multiple tissues and cell-types generated for the sheep gene expression atlas using mRNA- Seq (Clark et al. 2017).

**Figure 9.**
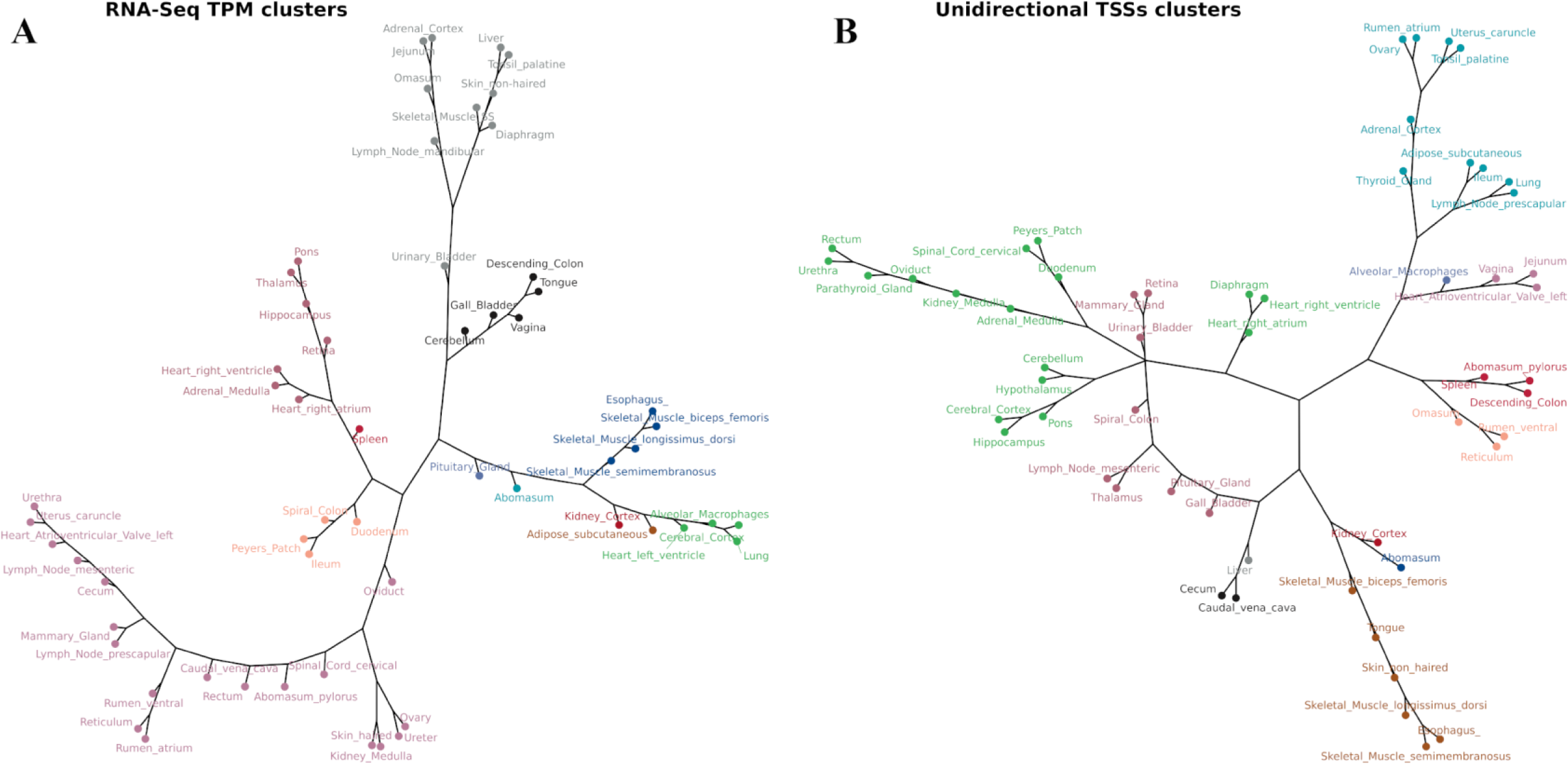
The network analysis of tissue TSS and gene expression profiles in 52 matched samples from Benz1626. The clustering algorithm was based on MI distance of each tissue given the expressed A) mRNA-Seq transcript level TPM and B) CAGE tag clusters (TSSs).

### Comparative visualisation of the datasets

An interactive visualisation interface was developed in order to make these datasets accessible and useable for the livestock genomics community. The genomic browser incorporates the bp resolution hypomethylation data, the CTPM expression of TSS and TSS-Enhancer regions and the mRNA-Seq TPM expression at transcript level. These tracks are also overlaid using the coordinates provided by the TxDB objects for transcripts and gene models as shown in Figure 10. This form of overlaid view allows for confirmation of transcript expression and the exact coordinate of the corresponding TSS in each tissue. For validation purposes the promoter region should be under a hypo-methylated CpG island on the DNA track for a proportion of actively transcribed gene in each tissue. The detailed bigBED format tracks for all the tissues are available at https://trackhubregistry.org/search/view_trackhub/TW3SmXMBjGhbrAAjJGTU ; https://data.faang.org/api/fire_api/trackhubregistry/hub.txt.

**Figure 10.**
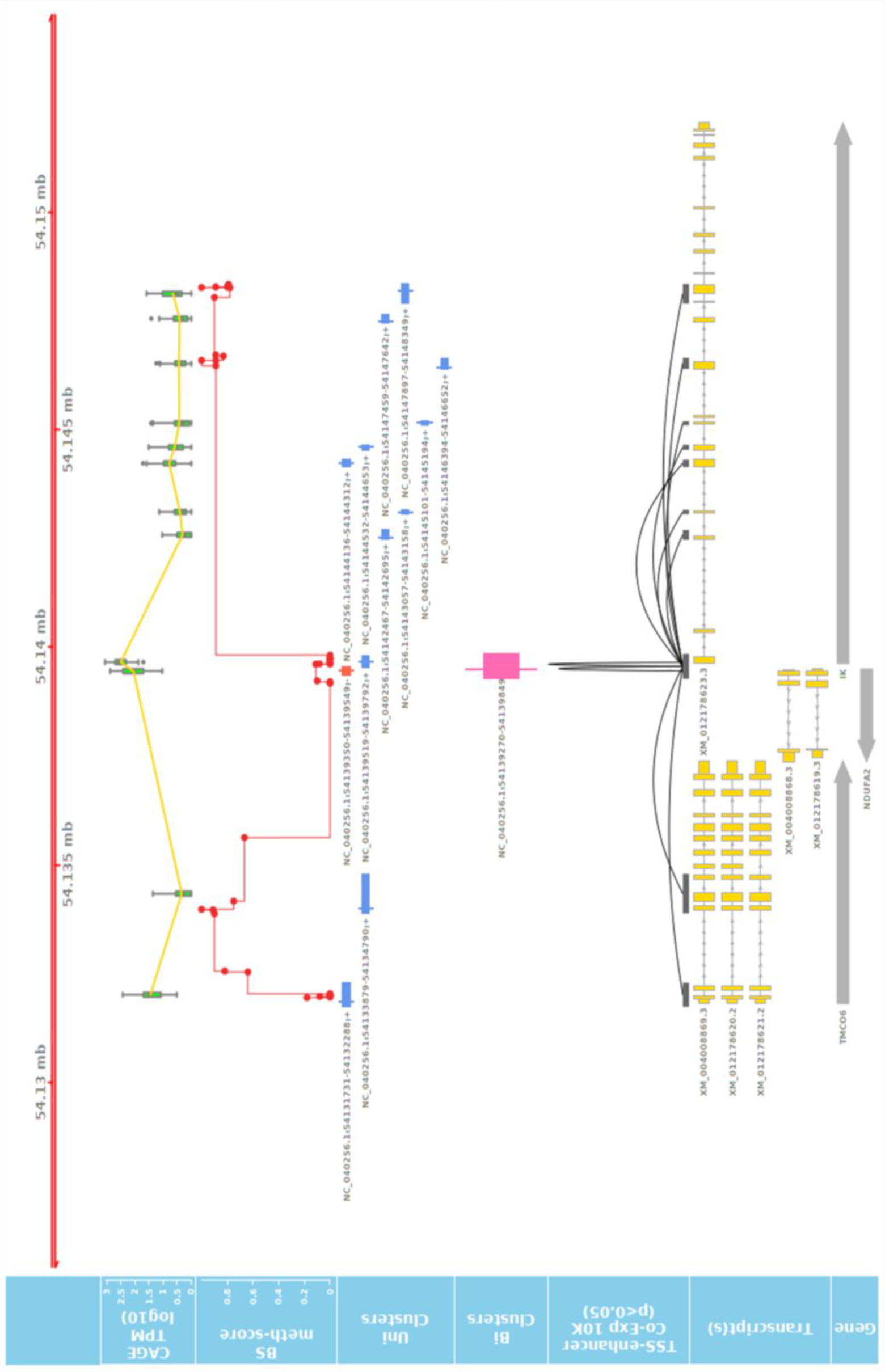
Long range correlation of single enhancer site with multiple promotors of several genes. The track shows the significant correlation of a leading/primary enhancer site highly co-expressed with several TSS sites of different genes in a relatively long coding frame (± 10,000Kb). The 3^rd^ track from the top also shows the level of methylation at CpG sites at DNA level of Benz2616 overlaying the same coordinates of the IK gene and ±10Kbp.

These visualisation tools were used to identify any co-expressed enhancers within the proximity of a TSS. We were able to identify 741 TSS-Enhancer clusters across the 56 tissues. An example of these bi-directional clusters is shown in Figure 10 as a pink line. The pairwise CTPM levels of co-expression of the bi-directional clusters and those of the uni-directional TSS clusters were compared using the Kendal correlation function in CAGEfightR (Thodberg and Sandelin, 2019). There were 5,383 significant co-expression pairs between uni-directional clusters (28,148) and bi- directional clusters (741). An example of a co-expressed TSS-enhancer is shown in Figure 10 as a black line connecting the significant start positions of the co-expression pairs.

The co-expression range of bi-directional clusters, in some cases, can span beyond the 10Kbp distance, as shown in the *IK* gene example (Figure 10). The expression of enhancer RNA (eRNA) with the promoter expression level of their target genes has been reported before (Tippens et al., 2018). This layer of annotation provides a foundation for enhancer target mapping in the sheep genome. The detailed list and annotated target transcripts of these co-expression clusters can be found in Supplementary File 2.

### FANTOM5 mammalian and avian CAGE TSS atlases

The FANTOM5 project also used CAGE to annotate TSSs in mammalian genomes (Andersson et al., 2014; Forrest et al., 2014b; Imada et al., 2020). The FANTOM5 data release contained putative TSSs for human, mouse, chicken, rhesus monkey and dog (https://fantom.gsc.riken.jp/5/data/). We performed a comparative analysis of the number of TSSs captured by these datasets with the CAGE dataset we generated for sheep (Table 2, Supplementary Table 6).

**Table 2:** Metrics comparison of CAGE atlases from 7 species. The total number of TSSs identified using CAGE methodology and the number of corresponding genes. Data for all the species other than sheep was accessed via the FANTOM5 data portal.

| Species | Genome | TSSs | Genes |
| --- | --- | --- | --- |
| Human | hg38 | 209,911 | 31,184 |
| Mouse | mm10 | 164,672 | 30,501 |
| Chicken | galGal5 | 32,015 | 7,759 |
| Sheep | Oar rambouillet v1.0 | 28,148 | 13,912 |
| Rhesus Monkey | rheMac8 | 25,869 | 8,047 |
| Dog | canFam3 | 23,147 | 5,288 |

The number of genes annotated by CAGE uni-directional TSS clusters in this study was greater than chicken, rhesus monkey and dog produced as part of FANTOM5 project, however TSS annotation for sheep was still consistently less robust than for murine and human genomes.

## Discussion

High-quality reference genomes are now available for many farmed animal species including domestic sheep (*Ovis aries*). The earlier draft genome sequence (Jiang et al., 2014) has been superseded by a more contiguous genome assembly (*Oar rambouillet v1.0* https://www.ncbi.nlm.nih.gov/assembly/GCF_002742125.1/). Annotation of this genome sequence, however, is currently limited to gene and transcript models. There is a lack of information on regulatory sequences and the complexity of the transcriptome is underestimated. For example, promoters and TSS are not well-annotated and alternative promoters and transcripts are poorly characterised. The overall aim of the Ovine FAANG project was to provide a comprehensive annotation of *Oar rambouillet v1.0*. To contribute to this aim we generated a high-resolution global annotation of transcription start sites (TSS) for sheep. After removal, of CAGE tags with < 10 read counts, 39.3% of TSS overlapped with 5’ ends of transcripts, as annotated previously by NCBI. A further 14.7% mapped to within 50bp of annotated promoter regions. Intersecting these predicted TSS regions with annotated promoter regions (±50bp) revealed 46% of the predicted TSS were ‘novel’ and previously un-annotated. Using whole genome bisulphite sequencing data from the same tissues we were able to determine that a proportion of these ‘novel’ TSS were hypo-methylated (32.2%) indicating that they are likely to be reproducible rather than ‘noise’. The number of NCBI transcript/gene models for which there was no associated CAGE tag cluster was relatively small (7%) when we removed the strict filtering criteria, indicating the usefulness of CAGE data for genome annotation. However, the ‘noisy’ nature of CAGE data, proportion of multi-mappers and duplicated reads, resulted in a considerable attrition of raw reads. We also chose to use strict filtration criteria, requiring the CAGE tags to be present in 2/3^rds^ of tissues. This resulted in a relatively modest number of high confidence CAGE clusters. This strict filtering could be relaxed for future analysis of the data. The global annotation of TSS in sheep we present will significantly enhance the annotation of gene models in the new ovine reference assembly (*Oar rambouillet v1.0)*.

The quality of the annotation of reference genomes for livestock species is improving rapidly with reductions in the cost of sequencing and generation of new datasets from multiple different functional assays (Giuffra and Tuggle, 2019). *Oar rambouillet v1.0* superseded the Texel reference assembly (*Oar_v3.1*), which was released in 2014 (Jiang et al., 2014). *Oar_v3.1* is still widely utilised by the sheep genomics community and the Ensembl annotation (https://www.ensembl.org/Ovis_aries/Info/Index) also includes sequence variation information. We compared how mapped CAGE tag clusters were distributed across genomic features in *Oar rambouillet v1.0* and *Oar_v3.1* (Jiang et al., 2014) and found that the proportion of CAGE tag clusters mapping to promoter regions was greater for *Oar rambouillet v1.0* (39%) than *Oar_v3.1* (15%). This may be because *Oar_v3.1* was built using short read technology (Jiang et al., 2014), which had a significant bias to GC rich regions, and therefore did not robustly capture the 5’ ends of many genes (Chen et al., 2013). In comparison, the *Oar rambouillet v1.0* assembly was generated using long read technology, that dramatically improves the ease of assembly resulting in increased contiguity (Contig N50: *Oar_v3.1* 0.07Mb and *Oar rambouillet v1.0* 2.57Mb). Other recent high quality reference genome assemblies for livestock, e.g. goat (Bickhart et al., 2017; Worley, 2017) and water buffalo (Low et al., 2019), have been built using long read sequencing technology in combination with optical mapping for scaffolding. Highly annotated genomes are powerful tools that can help us to understand the mechanisms underlying complex traits in livestock (Georges et al., 2018; Giuffra and Tuggle, 2019) and mitigate future challenges to food production (Rexroad et al., 2019). GWAS results, for example, can be integrated with functional annotation information to identify causal variants enriched in trait-linked tissues or cell types (reviewed in (Cano-Gamez and Trynka, 2020)). Using enrichment analysis (Finucane et al., 2018) showed that heritable disease associated variants from GWAS were enriched in enhancer regions in relevant tissues and cell types in humans. The TSS and TSS-enhancer clusters identified in this study could be utilised in a similar way for SNP enrichment analysis of GWAS variants in sheep. Using ChIP-Seq data (Naval-Sanchez et al., 2018) found that selective sweeps were significantly enriched for proximal regulatory elements to protein coding genes and genome features associated with active transcription. A high quality set of variants for sheep, generated using whole genome sequencing information for hundreds of animals across multiple breeds, is available through (Sheep Genomes Database, 2020). This dataset could be used to identify functional SNPs enriched in the TSS and TSS-enhancer clusters for multiple tissues and cell types that we have annotated in the *Oar rambouillet v1.0* assembly. High throughput functional screens using gene editing technologies, are now possible to validate these functional variants (reviewed in (Tait-Burkard et al., 2018)). New iPSC lines for livestock species also now offer the potential to do this in relevant cell types (Ogorevc et al., 2016).

Our high-resolution atlas of TSS complements other available large-scale RNA-Seq datasets for sheep e.g. (Clark et al., 2017). The analysis we present here includes tissues representing all major organ systems. However, we were unable to generate CAGE libraries for a small number of difficult to collect or problematic tissues, and as such may have missed transcripts specific to these tissues. We were also only able to generate CAGE libraries from one isolated cell type, alveolar macrophages. As demonstrated by the FANTOM5 (Forrest et al., 2014a) and ENCODE (Birney et al., 2007) and FragENCODE (Foissac et al., 2019) projects, including a diversity of immune cell types, in both activated and inactivated states, in future work would capture additional transcriptional diversity. New technologies, such as single cell sequencing, will allow annotation of cell-specific expressed and regulatory regions of the genome at unprecedented resolution (Papatheodorou et al., 2019). C1 CAGE now offers the opportunity to detect TSS and enhancer activity at single-cell resolution (Kouno et al., 2019).

We have also generated full-length transcript information using the Iso-Seq method, for a small subset of tissues from Benz2616. Integrating mRNA-Seq and Iso-Seq datasets has been used successfully to improve the annotation of the pig genome (Beiki et al., 2019). By merging the Iso-Seq data with the CAGE and mRNA-Seq datasets we will be able to measure differential transcript usage across tissues and improve the resolution of the *Oar rambouillet v1.0* transcriptome further. Our analysis indicated that while the vast majority of transcripts had one TSS, some genes had multiple putative TSS which could be validated with the additional resolution provided by the Iso-Seq data. As such the study we present here represents just the first step in demonstrating the power and utility of the different datasets generated for the Ovine FAANG project, which will provide one of the highest resolution annotations of transcript regulation and diversity in a livestock species to date.

## Acknowledgements

The authors would like to thank the Human Genome Sequencing Center staff for the *Oar rambouillet v1.0* reference genome assembly and for long-read and short read mRNA and miRNA sequencing. HGSC contributors include the genome assembly and analysis team of Y. Liu, R.A. Harris, X. Qin, led by K. Worley and production team members including M-C Gingras and L. Perez for RNA and DNA preparation, V. Vee, Y. Han. V. Korchina, S. Dugan-Perez for sequencing, Q. Meng, H. Doddapaneni, M. Wang for library production, the system support and LIMs teams, D. M. Muzny the HGSC Director of Operations, R. A. Gibbs the HGSC Director. We thank S. Sullivan and I. Liachko of Phase Genomics for PGA genome scaffolding.

We would also like to thank William Thompson for isolation of RNA and Jacky Carnahan for isolation of DNA at USMARC. The authors are grateful to Lucas Lefevre, Rachel Young and Heather Finlayson for performing the initial optimisation to establish the CAGE protocol at the Roslin Institute and Sara Clohisey and Lee Murphy for advice on data analysis and assistance in establishing the protocol at the Edinburgh Clinical Research Facility. CAGE libraries were sequenced at the Centre for Genomic Research, University of Liverpool. The authors are also grateful for the support of the FAANG Data Coordination Centre (http://data.faang.org) in the upload and archiving of the sample data and metadata, and hosting of the genome tracks.

Furthermore, we would like to thank the tissue collection design team; Noelle Cockett and Kim Worley, who coordinated the team, James Kijas, Brian Dalrymple, Tracy Hadfield, Kara Thornton, Tom Baldwin, Shuna Jones, Bob Lee, Sue Hauver, Christy Kelley, Jessica Eisenhauer, Mike Heaton, William Thompson, Timothy Smith, Stephen White, Michelle Mousel, Alisha Massa; Brian Sayre and Brenda Murdoch who all contributed in planning and designing the collection. Additionally, we would like to thank all those that contributed to the FAANG tissue collection at USU: Noelle Cockett, Tracy Hadfield, Tom Baldwin, Rusty Stott, Arnaud Van Wettere, Holly Mason, Jaqueline LaRose, Dave Forrester, Corey Wareham, Sarah Behunin, Kara Thornton, Gordon Hullinger (VMES), Alisha Massa, Maria Herndon, Brenda Murdoch, Brian Sayre, Caylee Birge, Codie Durfee, Michelle R. Mousel, Rachael Christianson, Nicole Ineck, Angie Robinson, Dallin Wengert, Kerry Rood, Erica Moscoso, Rickie Warr, Dustin Kinney, Abbey Benninghoff, Sumira Phatak, Kevin Contreras, Braden Abercrombie, Misha Regouski.

This manuscript has been released as a pre-print at bioRxiv, (Salavati et al., 2020).

## Ethics Statement

All protocols were approved by the animal care and use in accordance with Utah State University. IACUC approval: #2826, expiration date 21^st^ of February 2021.

## Data Availability

All the raw sequence data and analysis BAM files for this study are publicly available via the OAR_USU_Benz2616 NCBI BioProject: https://www.ncbi.nlm.nih.gov/bioproject/PRJNA414087 and via the European Nucleotide Archive (ENA): https://www.ebi.ac.uk/ena/browser/view/PRJEB34864 (CAGE), https://www.ebi.ac.uk/ena/data/view/PRJEB35292 (mRNA-Seq) and http://www.ebi.ac.uk/ena/data/view/PRJEB39178 (WGBS). Details of all 100 samples collected from Benz 2616 are included in the BioSamples database under submission GSB-7268, group accession number SAMEG329607 (https://www.ebi.ac.uk/biosamples/samples/SAMEG329607). The datasets are accessible via the FAANG data portal and were submitted according to FAANG sample and experimental metadata requirements (Harrison et al., 2018). Oar rambouillet v1.0 is now available on the Ensembl Genome Browser http://www.ensembl.org/Ovis_aries_rambouillet/Info/Index.

## Author Contributions

RC and IG performed CAGE library, optimisation, preparation and sequencing. MS performed all bioinformatic and data analyses, with the exception of the WGBS data, which was analysed by AC. MS and AC generated the GViz tracks. TPLS coordinated generation of the mRNA-Seq data at US-MARC. KCW coordinated generation of the Oar Rambouillet v1.0 reference assembly and mRNA-Seq data at BCM. SMC coordinated the generation and analysis of the WGBS with AC. ELC and ALA coordinated the CAGE components of the study. NEC and KCW planned and coordinated the sample collection at USU. BMM is coordinator of the Ovine FAANG project with ALA, SNW, BPD, JWK, RB, NEC, SMC, KCW, ELC and TPLS who designed the overall project and acquired the funding to support the work. ELC wrote the manuscript with MS and AC. All authors contributed to editing and approved the final version of the manuscript.

## Code Availability

All the code base for the analytical pipeline in this study are available at https://msalavat@bitbucket.org/msalavat/rnaseqwrap_public.git for RNA-Seq analysis, https://msalavat@bitbucket.org/msalavat/cagewrap_public.git for the CAGE mapping, annotation and metrics pipeline and https://msalavat@bitbucket.org/caultona/wgbswrap_public.git for WGBS pipeline.

## Members of the Ovine FAANG Project Consortium (listed by institution)

Brenda Murdoch (University of Idaho)

Kimberly M Davenport (University of Idaho)

Stephen White (USDA, ARS, Washington State University)

Michelle Mousel (USDA, ARS ADRC)

Alisha Massa (Washington State University)

Kim Worley (Baylor College of Medicine)

Alan Archibald (The Roslin Institute, University of Edinburgh)

Emily Clark (The Roslin Institute, University of Edinburgh)

Brian Dalrymple (University of Western Australia)

James Kijas (CSIRO)

Shannon Clarke (AgResearch) Rudiger Brauning (AgResearch)

Timothy Smith (USDA, ARS MARC)

Tracey Hadfield (Utah State University)

Noelle Cockett (Utah State University)

## Supplemental Material

**Supplementary Figure S1.**
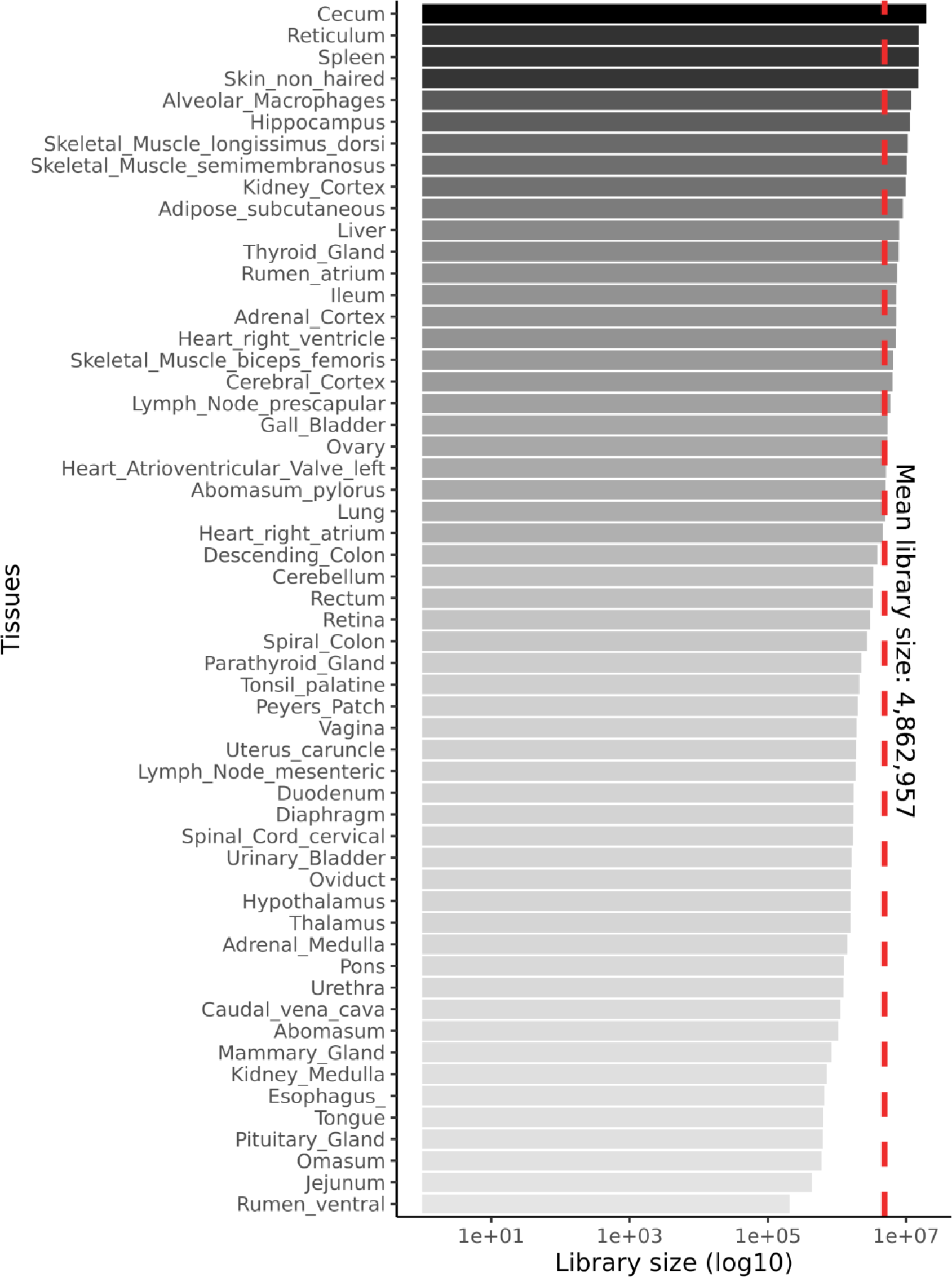
CAGE library size for each of the 56 tissues analysed.

**Supplementary Figure S2.**
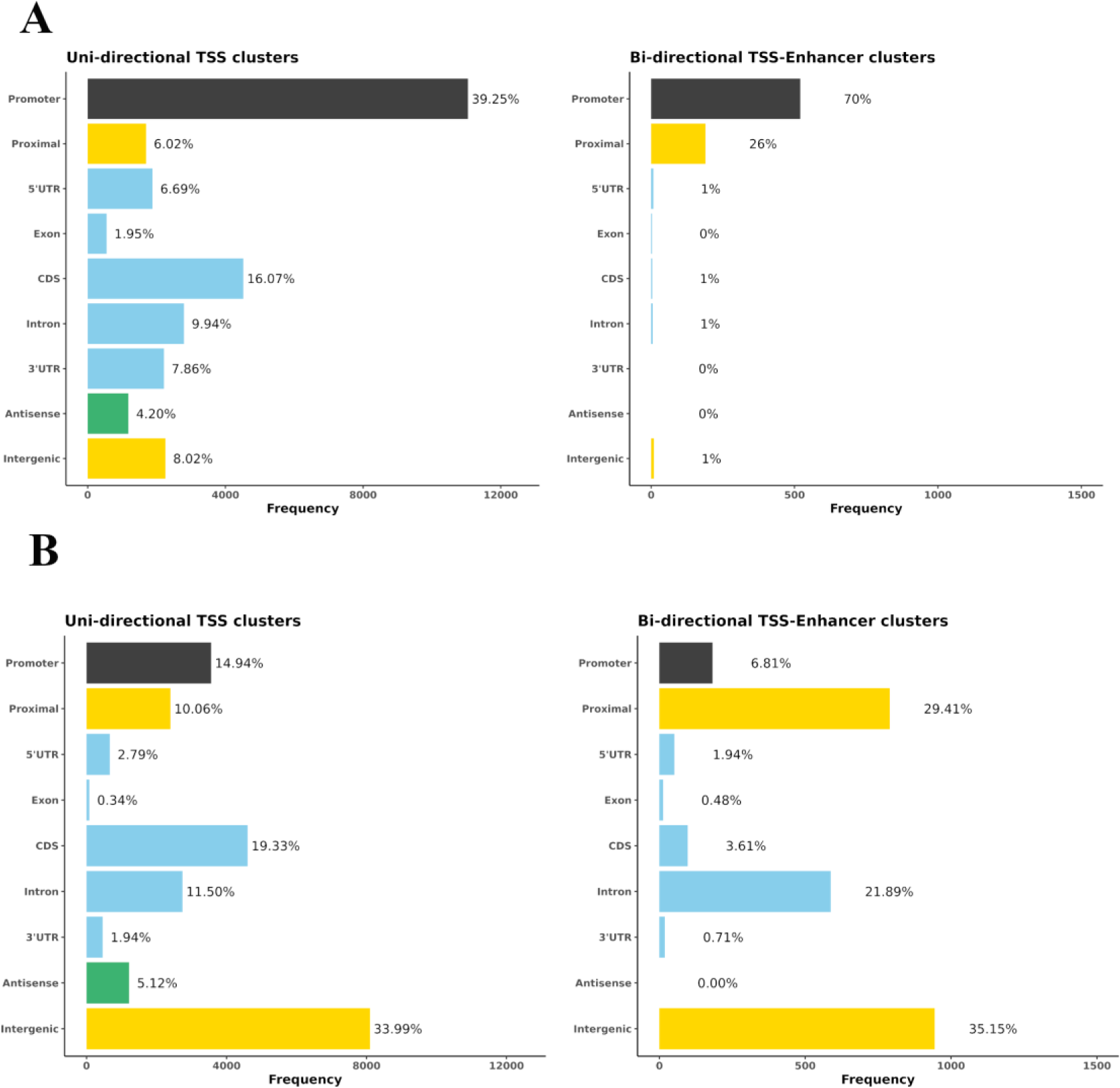
The percentage of CAGE tags mapped to each genomic region for Oar rambouillet v1.0 (A) and Oar_v3.1 (B) reference genome assemblies. The counts were averaged across tissues prior to annotation.

**Supplementary Figure S3A.**
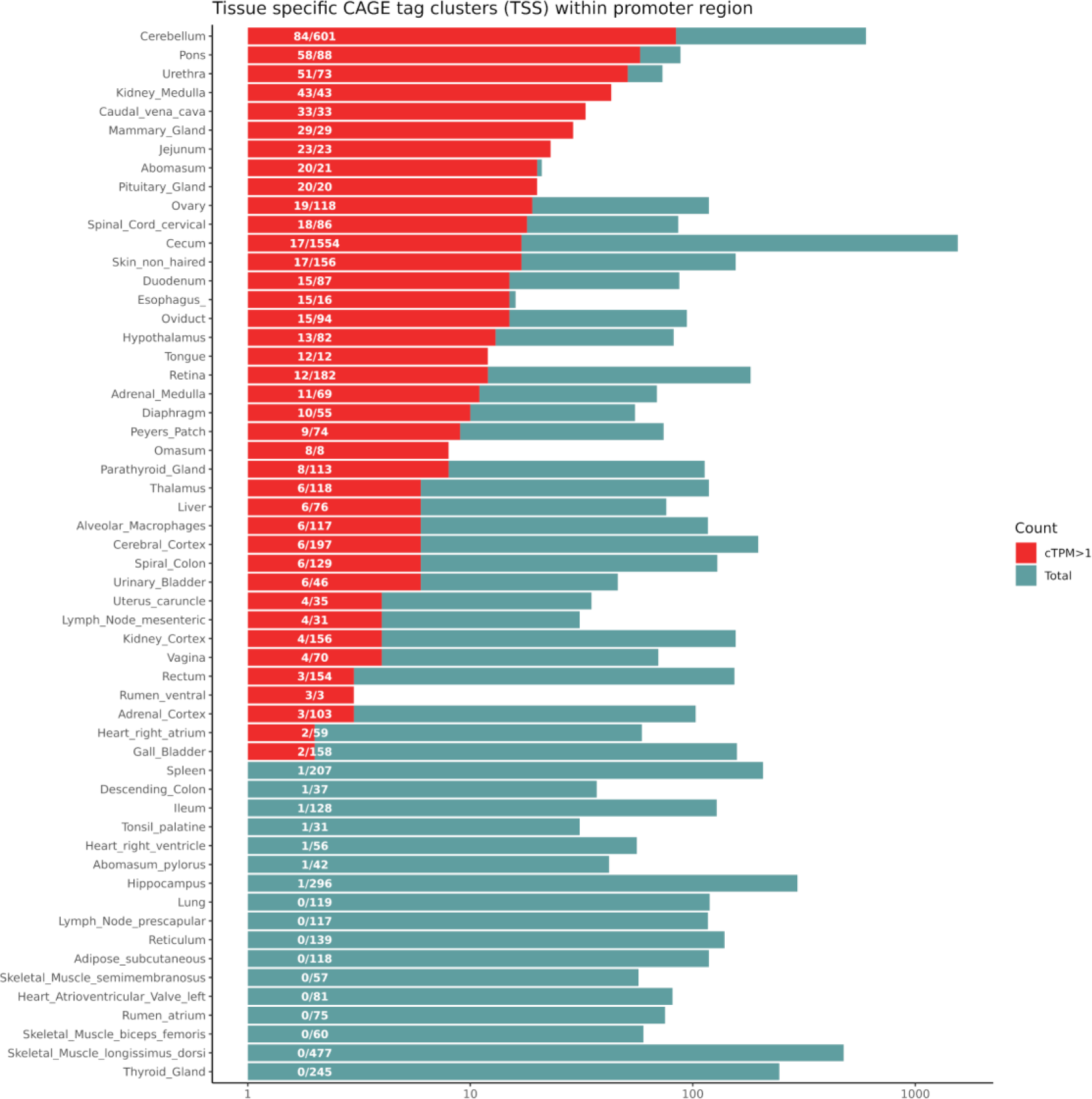
The distribution of tissue specific TSS in 56 tissues of Benz2616. The bar shows the count of tissue specific TSS in each tissue with the proportion being expressed with CTPM >1 coloured in red.

**Supplementary Figure S3B.**
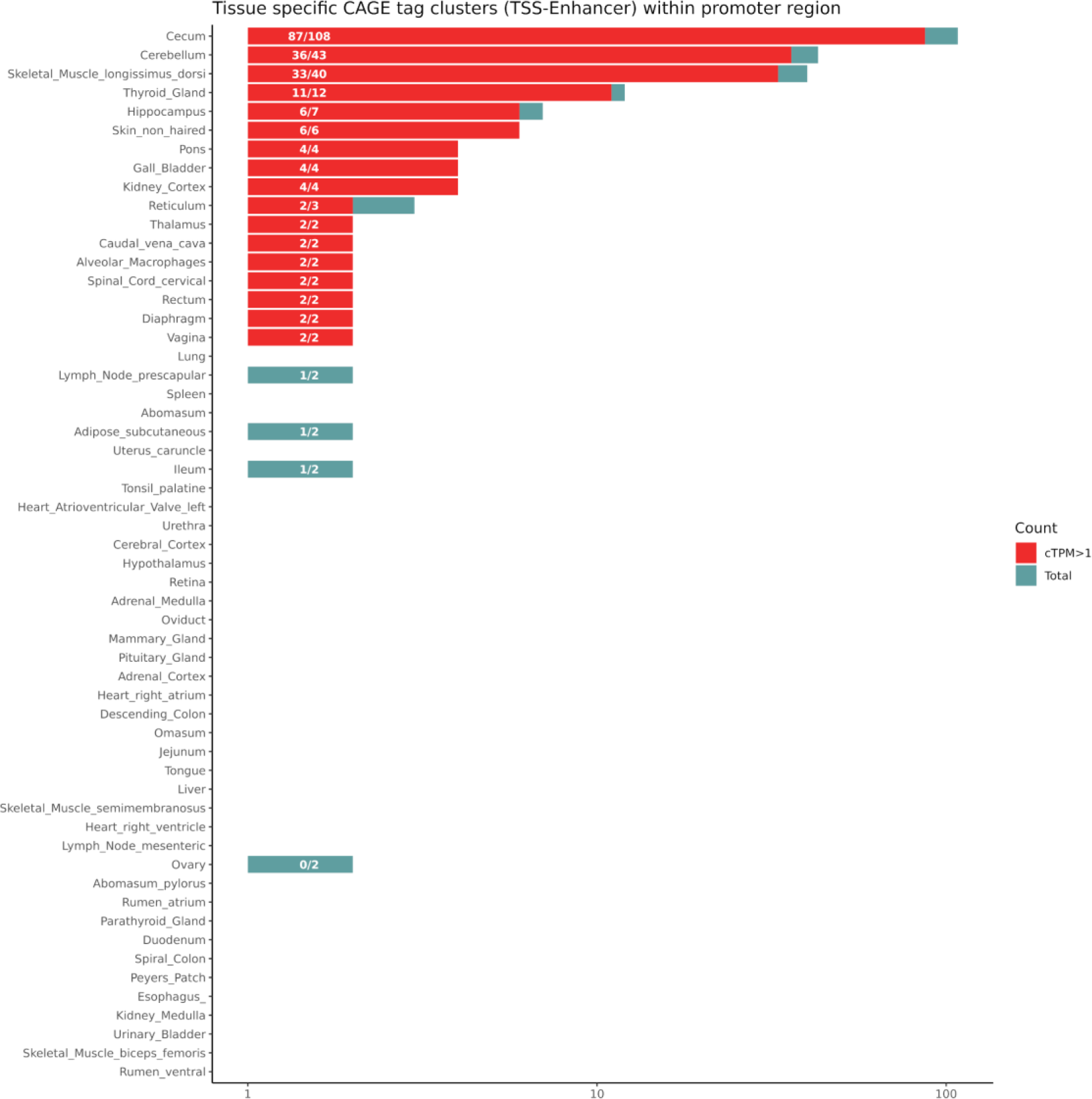
The distribution of tissue specific TSS-Enhancers across the 56 tissues from Benz2616. The bars show the count of tissue specific TSS in each tissue with the proportion being expressed with CTPM >1 coloured in red.

**Supplementary Figure S4.**
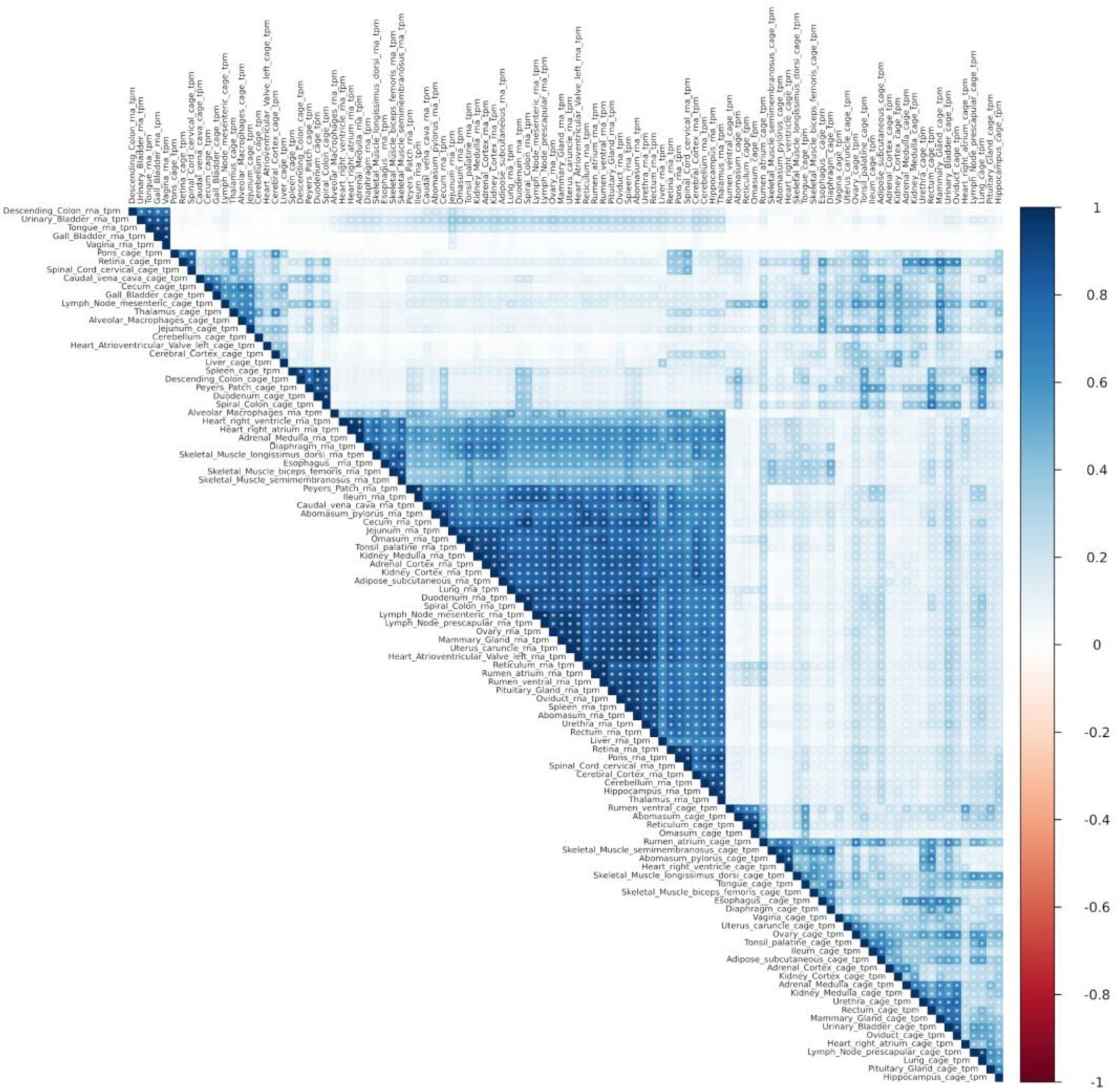
Heatmap of mRNA-Seq and CAGE expression profiles (TPM and CTPM). The correlation was calculated over 52 matched tissues and 5732 transcripts-TSS expressed in all tissues.

**Supplementary Figure S5.**
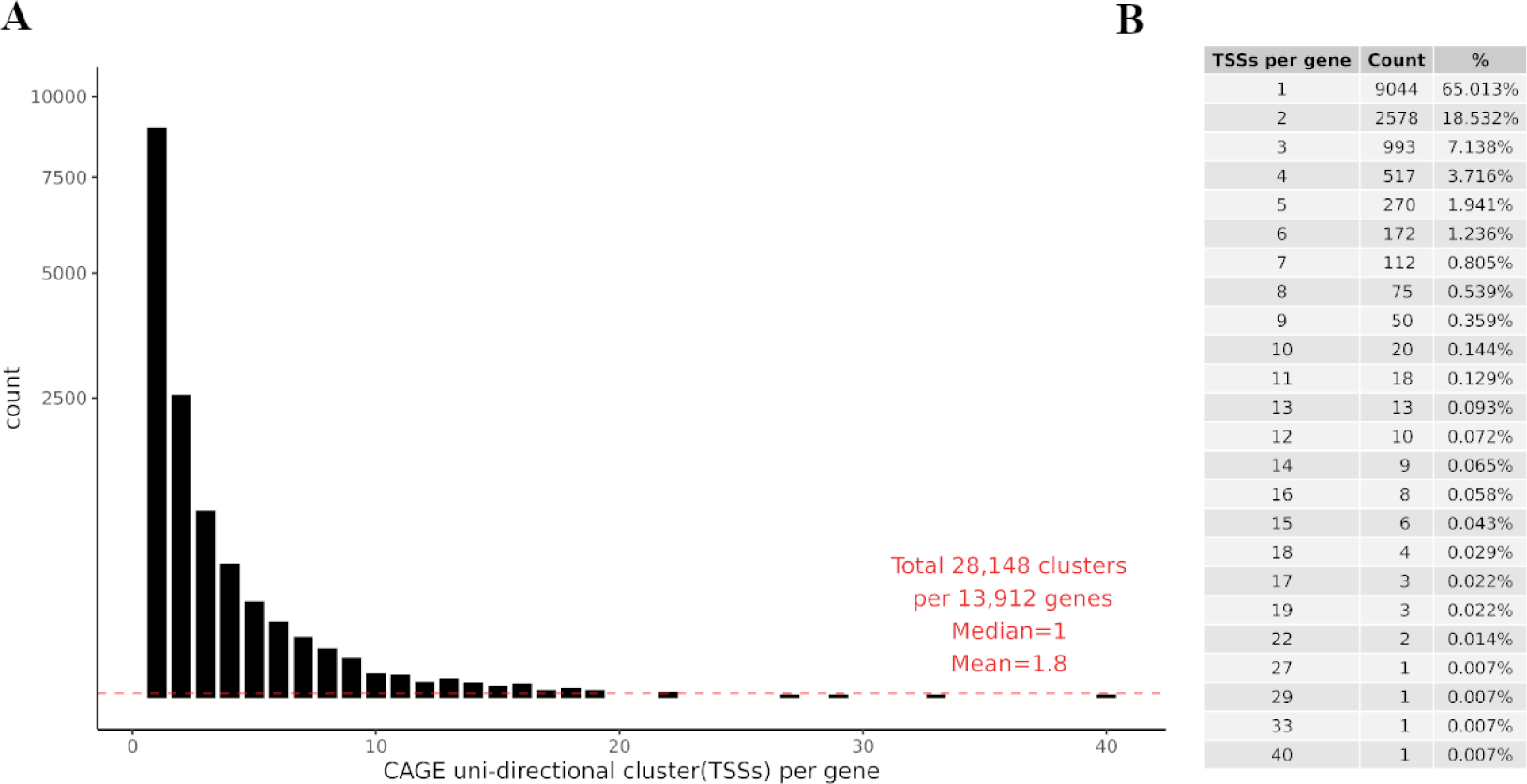
Distribution of uni-directional CAGE TSS clusters per annotated gene. A) the histogram of the TSS cluster per gene B) Detailed table of TSS per gene data underlying the histogram and percentage per total TSS clusters.

Supplementary Table 1. Details of 5’ linker barcodes and pool ID assigned to each tissue sample.

Supplementary Table 2. Percentage of tissue-specific CAGE tags mapping to genomic features.

Supplementary Table .3. Summary of WGBS sequencing and mapping results.

Supplementary Table 4. Comparison of mRNA-Seq dataset with matched tissues within the CAGE dataset with regards to tissue support criteria.

Supplementary Table 5. Summary of minimum tissue support calculations for TSSs per gene in each scenario. Tissue support thresholds of 1, 5, 18, 28 & 37 tissues out of total 56 were analysed.

Supplementary Table 6. Comparison of this study with other CAGE datasets available as part of the Fantom5 consortium data release.

Supplementary File 1. Section 1: Attrition of raw reads at each stage of the analysis pipeline, rationale for selecting the 2/3^rds^ representation threshold for mapped CAGE tags and clustering metrics. Sections 2 and 3: detailed comparison of mapping of the CAGE tags to the two reference assemblies Oar_v3.1 and Oar rambouillet v1.0 and analysis workflow.

Supplementary File 2. Expression data frames from Uni-, Bi-directional, long range linked co-expression clustering and transcript level mRNA-Seq from all 56 tissues (2/3^rds^ representation rule applied).

