## Supplementary material for "Global analysis of transcription start sites in the new ovine reference genome (*Oar rambouillet v1.0*)": Supp File 1

**Supplementary File 1:**

**Section 1**

**1.1 The overall input and output of the CAGE analysis pipeline -attrition of raw reads:**

This metric table below shows the attrition of reads from the raw fastq reads to mapped BAM format and ultimately the raw tag count. The clustered TSS with different levels of cTPM expression are summarised in the figures below. From an average 7.3 million fastq reads after trimming and mapping to the reference genome, 4.8 million uniquely mapped reads were retained (33% loss). These high quality mapped reads were then decomposed to bp resolution using bedtools, resulting in over 116 million tags per tissues on average. Across 56 tissues overall 5.4 million TSS clusters were identified, the majority of which had a cTPM between 0-1. On average 358k clusters (across 56 tissues) had expression levels of approximately 1 cTPM and only 8219 on average were expressed >10 cTPM when each tissue was analysed separately. As previously described the cTPM was calculated again after filtering these 5.4 million clusters using the 2/3^rd^ representation threshold to correct for the library size differences (please refer to the following table for the details).

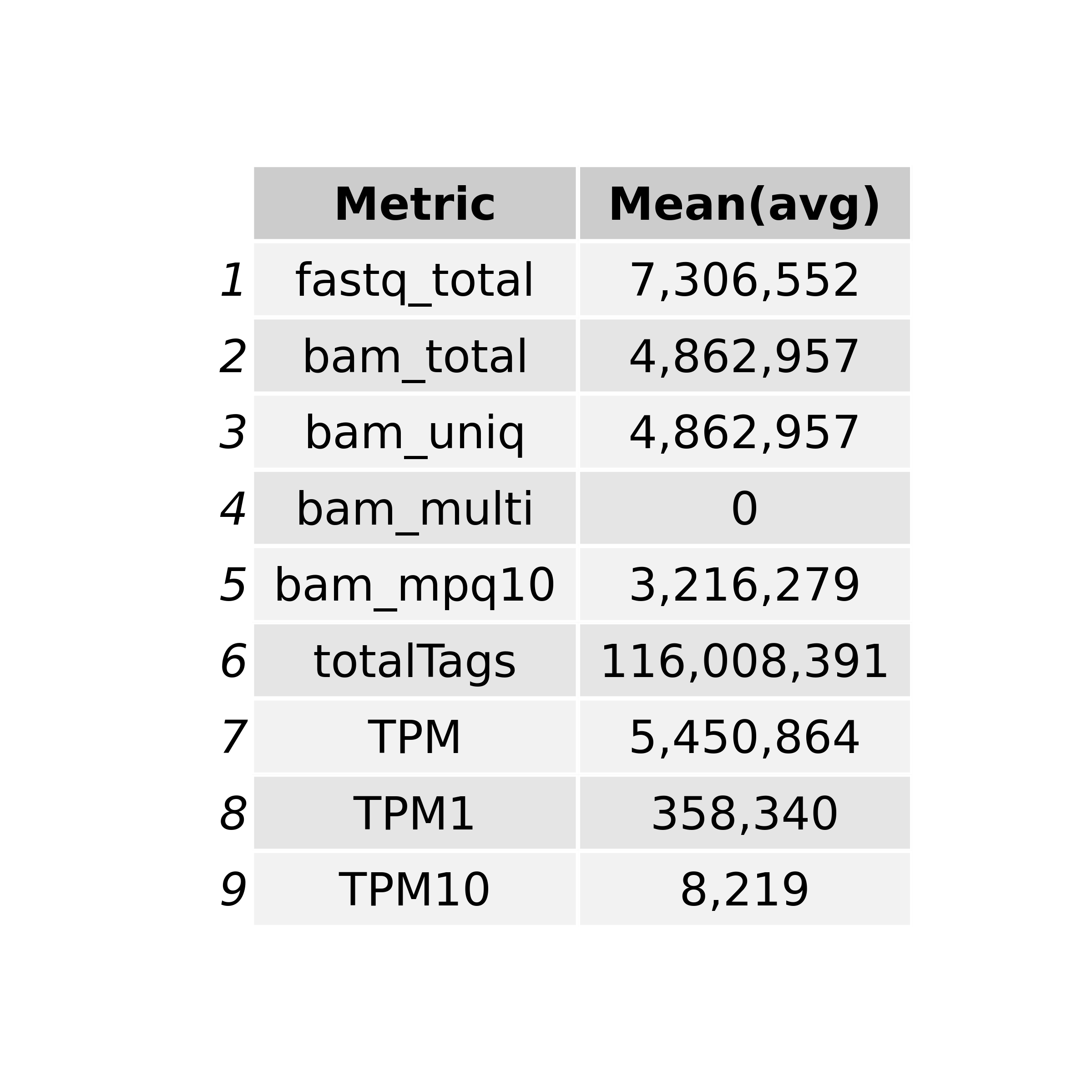

*Table 1.1 The overall averaged (across 56 tissue) metrics of the input and output of the CAGE analysis pipeline. Average values rounded up to the next digit.*

*fastq_total = Average number of the total reads in each fastq file; bam_total = Average number of the mapped reads in each bam file ; bam_uniq= Average number of the uniquely mapped reads in each bam file; bam_multi= Average number of multi-mapped reads in each bam file; bam_mpq10 = Average number of reads with mapping quality Phred score above 10; totalTags = Average number of the decomposed tags (bigWig +/-) from each bam file ; TPM = Total number of clustered TSS across 56 tissues with TPM > 0; TPM1 for >1 and TPM10 for >10*

**1.2 Logic for selecting a 2/3^rd^ representation threshold for mapped CAGE tag clusters for each gene shared across tissues from Benz2616**

The representation threshold for CAGE tag clusters that were shared across tissues for each gene was optimised using the steps described below. The optimisation range was compared using the total number of geneIDs (any NA IDs were removed) from the gff3 file. The TSS tag clusters that did not map to intronic or intergenic regions were separated into subsets. The untagged percentage represents geneIDs without any TSS or TSS-Enhancer assignment in the dataset. Three representation thresholds were compared 1 tissue, 1/3rd of tissues, ½ of tissues and 2/3rds of tissues. The threshold with the lowest proportion of ‘untagged’ was considered optimal to minimise ‘noise’ caused by un-annotated probably spurious TSS tag clusters in downstream analysis. According to this criteria we selected the 2/3rds threshold, which although stringent, gave us only the highest confidence TSS tag clusters, associated with widely expressed genes and widely used promotors, for the analysis of the dataset presented here. Providing a blind test, applying the same representation criteria to the mRNA-Seq dataset (n=58 tissues polyA enriched) created a similar pattern of reduction in transcripts being expressed (TPM 1 or TPM 10) and present in multiple tissues

The R script for the comparison in gene and transcript IDs is shown below:

*TSSs_full %>%*

*subset(txType != "intergenic") %>%*

*subset(txType != "intron") %>%*

*subset(geneID != "NA") %>%*

*subset(support > S) %>%*

*rowRanges() %>% as.data.frame() %>% dplyr::select(geneID) %>% unique() %>% count(geneID) %>% lengths() %>% .['n']*

| Biological support(S) | TSS clusters | tagged genes | total genes | untagged |
| --- | --- | --- | --- | --- |
| 1 tissue | 369,369 | 28,668 | 30,862 | 7.1% |
| 5 tissues | 196,349 | 22,938 | 30,862 | 25.6% |
| 18 tissues (1/3^rd^) | 62,735 | 18,005 | 30,862 | 41.6% |
| 28 tissues (1/2^nd^) | 35,806 | 15,727 | 30,862 | 49.0% |
| 37 tissues (2/3^rd^) | 24,105 | 13,912 | 30,862 | 54.9% |
| All tissues | 3,037 | 2,949 | 30,862 | 90.1% |

*TSSs_full %>%*

*subset(txType != "intergenic") %>%*

*subset(txType != "intron") %>%*

*subset(txID != "NA") %>%*

*subset(support > S) %>%*

*rowRanges() %>% as.data.frame() %>% dplyr::select(txID) %>% unique() %>% count(txID) %>% lengths() %>% .['n']*

| Biological support(S) | TSS clusters | tagged tx | total tx | untagged |
| --- | --- | --- | --- | --- |
| 1 tissue | 369,369 | 53,507 | 56,308 | 5.0% |
| 5 tissues | 196,349 | 46,574 | 56,308 | 17.3% |
| 18 tissues (1/3^rd^) | 62,735 | 39,458 | 56,308 | 29.9% |
| 28 tissues (1/2^nd^) | 35,806 | 35,492 | 56,308 | 37.0% |
| 37 tissues (2/3^rd^) | 24,105 | 31,113 | 56,308 | 44.7% |
| All tissues | 3,037 | 6,946 | 56,308 | 87.7% |

The breakdown of minimum support comparison and number of putative TSS clusters per gene is provided in the *min_support_comparison.xlsx* spreadsheet.

Providing a blind test, applying the same representation criteria to the mRNA-Seq dataset (n=58 tissues polyA enriched) created similar pattern of reduction in transcripts being expressed (TPM 1 or TPM 10) and present in multiple tissues:

| RNA-Seq captured transcripts | Tagged_tx | Total tx | untagged |
| --- | --- | --- | --- |
| Total (TPM >1) | 50,822 | 56,308 | 9.7% |
| TPM >10 | 32,852 | 56,308 | 41.7% |
| TPM >10 Tissue support 1 | 26,549 | 56,308 | 52.9% |
| TPM >10 Tissue support 5 | 16,936 | 56,308 | 69.9% |
| TPM >10 Tissue support 1/3^rd^ (19) | 8,427 | 56,308 | 85.0% |
| TPM >10 Tissue support 1/2^nd^ (29) | 5,914 | 56,308 | 89.5% |
| TPM >10 Tissue support 2/3^rd^ (38) | 3,908 | 56,308 | 93.1% |

The complete tissue by tissue break down of captured transcripts can be found in Supplementary Table 4.

**1.3 The relationship between pre vs post clustering metrics in the CAGE analysis pipeline (viz. uni-directional clustering algorithm of CAGEfightR).**

The regressed relationship between total tags recovered in each tissue and number of TSS clusters (TPM>10) was analysed using a series of linear and polynomial equations as shown in the figure below. The total tag count in each tissue was compared with Total reads (fastq; A), Total uniquely mapped read (BAM;B) and Total uniquely mapped reads with mapping quality >10 (BAM;C). The same comparison was also carried out for the final TSS cluster count (TPM>10) for each tissue in section D,E, and F respectively. The relationship between input reads and metrics in section A,B&C are positive with R^2^ ranging from 0.68 to 0.98 in linear and 0.88 to 0.98 using 3^rd^ degree polynomial regression. However; post clustering algorithm of CAGEfightR (Uni-directional) this positive relationship is absent as shown in sections D,E&F with R^2^ ranging from 0.027-0.023.

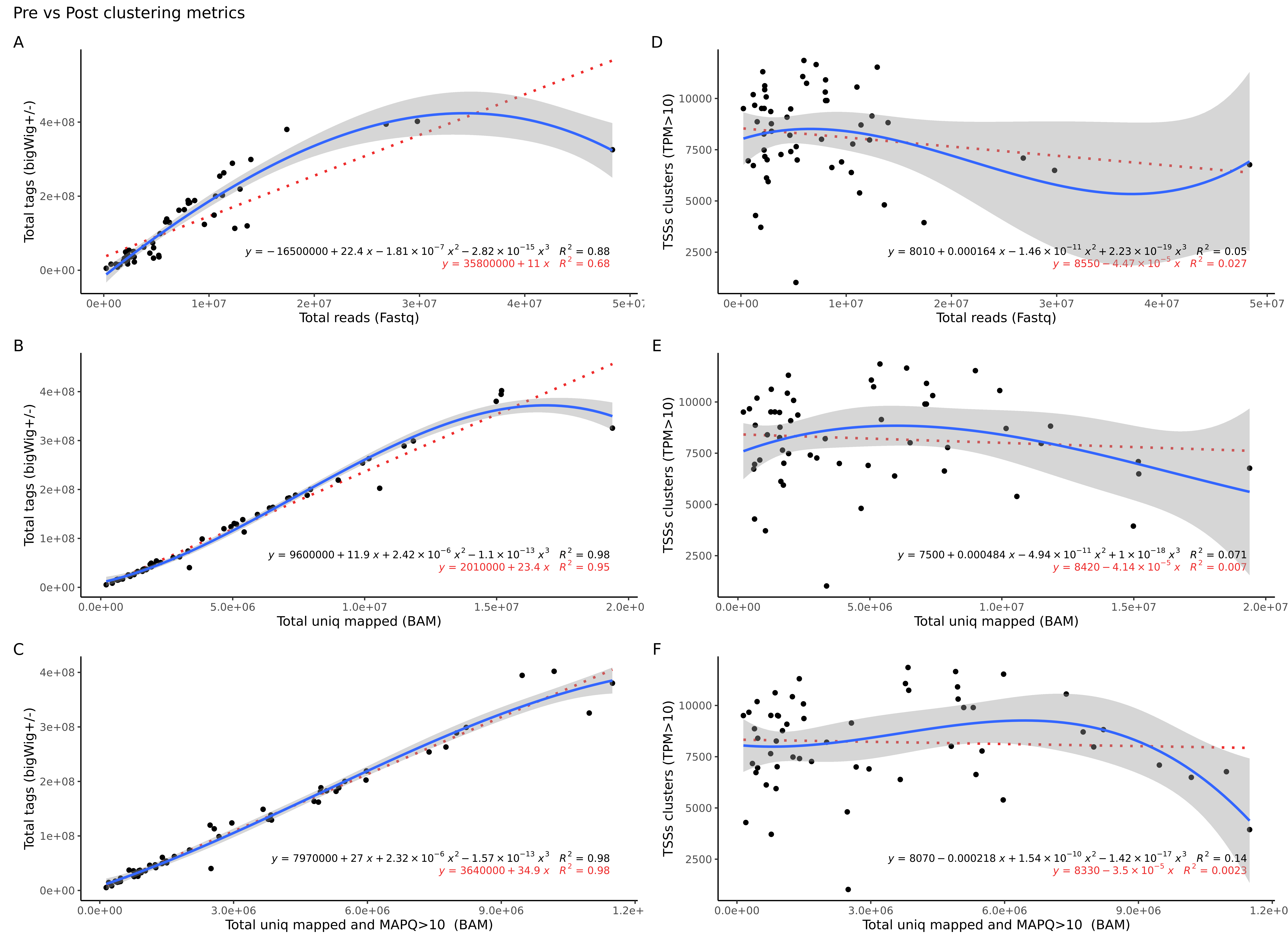

**Section 2.** The GFF file was downloaded for each reference assembly from NCBI v99 and Ensembl v99. The GFF files were compared for the number of features available in each category. This document provides a detailed description of the comparison between the two assemblies.

**2.1 The GFF3 comparison (gffcompare) metrics**

Summary for dataset: Ovis_aries.Oar_v3.1.99.gff3

Query mRNAs : 29079 in	26899 loci (22841 multi-exon transcripts)

(**2047 multi-transcript loci, ~1.1 transcripts per locus**)

Summary for dataset: GCF_002742125.1_Oar_rambouillet_v1.0_genomic.gff

Query mRNAs : 57320 in	33405 loci (47559 multi-exon transcripts)

(**8912 multi-transcript loci, ~1.7 transcripts per locus**)

Total union super-loci across all input datasets: 60304

(**10842 multi-transcript, ~1.4 transcripts per locus**)

86399 out of 86399 consensus transcripts written in gffcmp.combined.gtf (0 discarded as redundant)

**2.2 Feature type comparison (counts)**

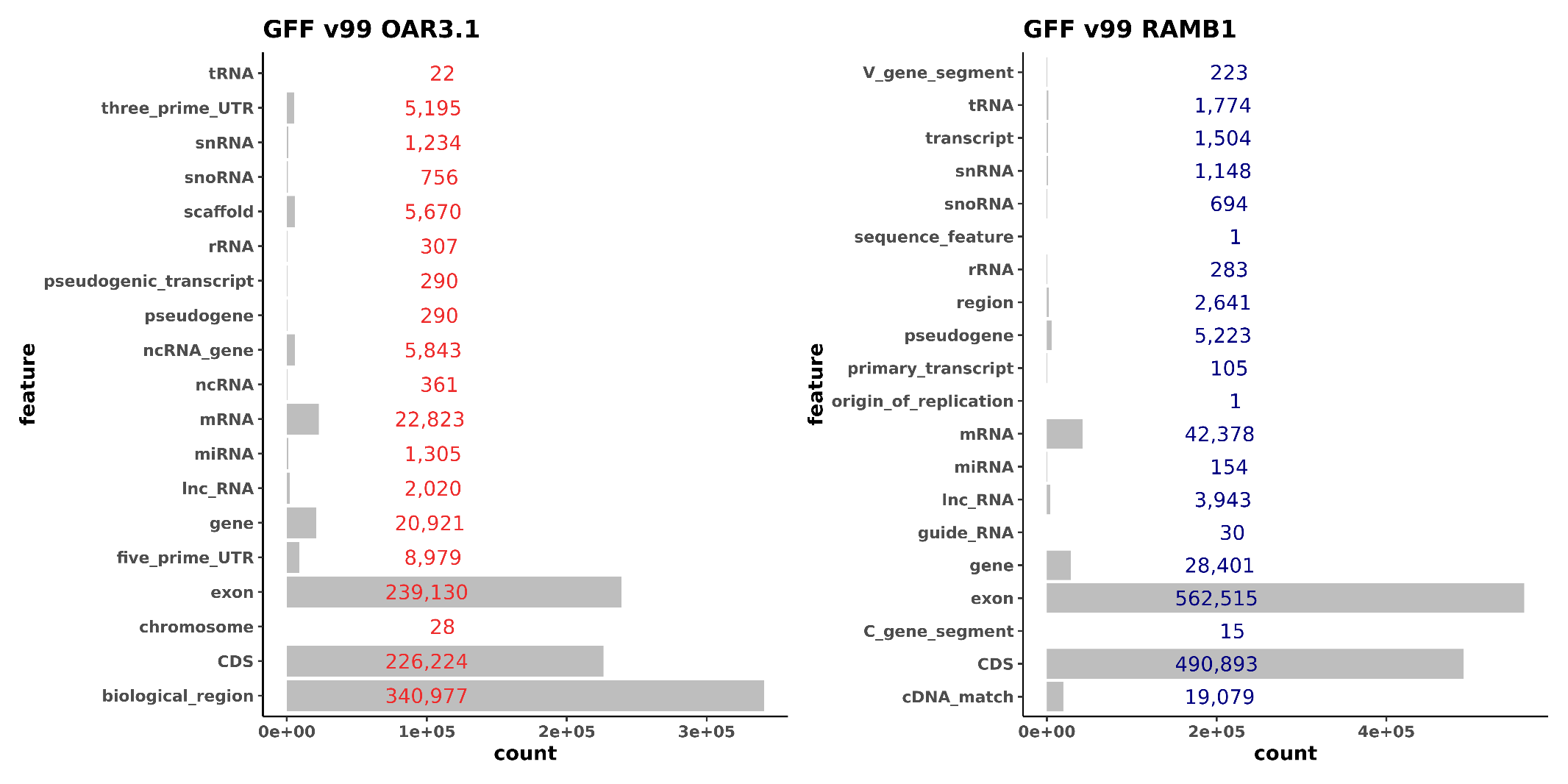

BSgnome and TxDB objects were generated from the GFF files according to the instructions available at <https://www.bioconductor.org/packages/devel/bioc/vignettes/BSgenome/inst/doc/BSgenomeForge.pdf>, <https://bioconductor.org/packages/release/bioc/vignettes/GenomicFeatures/inst/doc/GenomicFeatures.pdf>.

**2.3 TxDB object made from Oar_rambouillet_v1.0**

```
txdb
TxDb object:
# Db type: TxDb
# Supporting package: GenomicFeatures
# Data source: NCBI
# Organism: Ovis aries
# Taxonomy ID: 9940
# miRBase build ID: NA
# Genome: NA
# transcript_nrow: 57112
# exon_nrow: 561939
# cds_nrow: 490301
# Db created by: GenomicFeatures package from Bioconductor
# Creation time: 2020-05-28 13:23:49 +0100 (Thu, 28 May 2020)
# GenomicFeatures version at creation time: 1.36.4
# RSQLite version at creation time: 2.2.0
# DBSCHEMAVERSION: 1.2


promoters(txdb)
GRanges object with 56308 ranges and 2 metadata columns:
                                  seqnames        ranges strand |     tx_id                      tx_name
                                     <Rle>     <IRanges>  <Rle> | <integer>                  <character>
                  LOC101102048 NC_040252.1   19151-21350      + |         1                 LOC101102048
                XR_003588700.1 NC_040252.1   36177-38376      + |         2               XR_003588700.1
                XM_027964169.1 NC_040252.1 116194-118393      + |         3               XM_027964169.1
                XM_004003428.4 NC_040252.1 219799-221998      + |         4               XM_004003428.4
                XM_027966775.1 NC_040252.1 219808-222007      + |         5               XM_027966775.1
                           ...         ...           ...    ... .       ...                          ...
    rna-NC_001941.1:5263..5330 NC_001941.1     5131-7330      - |     57108   rna-NC_001941.1:5263..5330
    rna-NC_001941.1:6874..6944 NC_001941.1     6745-8944      - |     57109   rna-NC_001941.1:6874..6944
                           ND6 NC_001941.1   13886-16085      - |     57110                          ND6
  rna-NC_001941.1:14086..14154 NC_001941.1   13955-16154      - |     57111 rna-NC_001941.1:14086..14154
  rna-NC_001941.1:15371..15436 NC_001941.1   15237-17436      - |     57112 rna-NC_001941.1:15371..15436
```

**2.4 TxDB object made from Oar_v3.1**

```
txdb2
TxDb object:
# Db type: TxDb
# Supporting package: GenomicFeatures
# Data source: Ensembl
# Organism: Ovis aries
# Taxonomy ID: 9940
# miRBase build ID: NA
# Genome: NA
# transcript_nrow: 29118
# exon_nrow: 218775
# cds_nrow: 226224
# Db created by: GenomicFeatures package from Bioconductor
# Creation time: 2020-05-29 11:37:19 +0100 (Fri, 29 May 2020)
# GenomicFeatures version at creation time: 1.36.4
# RSQLite version at creation time: 2.2.0
# DBSCHEMAVERSION: 1.2

promoters(txdb2)
GRanges object with 28294 ranges and 2 metadata columns:
                                seqnames              ranges strand |     tx_id                       tx_name
                                   <Rle>           <IRanges>  <Rle> | <integer>                   <character>
                                       1         47751-49950      + |         1                          <NA>
                      PDCD1-201        1         77926-80125      + |         2                     PDCD1-201
  transcript:ENSOART00000027409        1         89112-91311      + |         3 transcript:ENSOART00000027409
                      DTYMK-201        1       184410-186609      + |         4                     DTYMK-201
                      THAP4-201        1       213759-215958      + |         5                     THAP4-201
                            ...      ...                 ...    ... .       ...                           ...
                                       X 134316336-134318535      - |     29114                          <NA>
                                       X 133935149-133937348      - |     29115                          <NA>
                                       X 134677341-134679540      - |     29116                          <NA>
                                       X 134842995-134845194      - |     29117                          <NA>
                     PABPC5-201        X 135116050-135118249      - |     29118                    PABPC5-201
```

**Section 3.** CAGE raw data was mapped against each assembly as previously described and subjected to the same CAGEfightR analysis workflow:

**3.1 Mapped against Oar_rambouillet_v1.0**

> TSSs

class: RangedSummarizedExperiment

dim: **28148** 56

> BCs

class: RangedSummarizedExperiment

dim: 741 56

**3.2 Mapped against Oar_v3.1**

> TSSs2

class: RangedSummarizedExperiment

dim: **23829** 56

>BCs2

class: RangedSummarizedExperiment

dim: **1121** 56

After application of the 2/3^rd^ representation threshold for the 56 tissues the supported CAGE tag clusters (i.e ‘Features’ as they are referred to in the CAGEfightR package) and unique annotation counts were generated as follows:

**3.3 Unidirectional TSS features**

AssignGeneID for Oar_rambouillet_v1.0 (Total = 28,148)

Features overlapping genes: 87.74 %

Number of unique genes: 13868

AssignTxID for Oar_rambouillet_v1.0 (Total = 28,148)

Features overlapping transcripts: 88.31 %

Number of unique transcripts: 31729

AssignGeneID for Oar_v3.1 (Total = 23,829)

Features overlapping genes: 49.1 %

Number of unique genes: 6549

AssignTxID for Oar_v3.1 (Total = 23,829)

Features overlapping transcripts: 65.41 %

Number of unique transcripts: 9914

**3.4 Bi-directional TSS-Enhancer features**

AssignGeneID for Oar_rambouillet_v1.0 (Total = 741)

Features overlapping genes: 87.74 %

Number of unique genes: 2598

AssignTxID for Oar_rambouillet_v1.0 (Total = 741)

Features overlapping transcripts: 88.92 %

Number of unique transcripts: 8075

AssignGeneID for Oar_v3.1 (Total = 1121)

Features overlapping genes: 54.58 %

Number of unique genes: 1371

AssignTxID for Oar_v3.1 (Total = 1121)

Features overlapping transcripts: 67.24 %

Number of unique transcripts: 2176
